# Sox8 and Sox9 regulate differentiation and nuclear positioning of retinal Müller glia

**DOI:** 10.64898/2026.01.12.698638

**Authors:** Nicole Pannullo, Debarpita Datta, Clayton P. Santiago, Jared A. Tangeman, Lizhi Jiang, Leighton H. Duncan, Seth Blackshaw

## Abstract

Temporal patterning of retinal progenitor cells governs the sequential generation of retinal cell types, with gliogenesis occurring late in development. Sox8 and Sox9, members of the SoxE transcription factor family, are highly expressed in late-stage retinal progenitor cells and mature Müller glia, yet their functional roles remain incompletely defined. Here we employed gain- and loss-of-function approaches, single-cell multiomic profiling, and injury models to investigate Sox8/9 function. Overexpression of *SOX8* and/or *SOX9* in early-stage retinal progenitor cells suppressed early-born cell fates and promoted photoreceptor generation, consistent with a role in late-stage temporal identity. Conversely, conditional deletion of *Sox8* and/or *Sox9* in late-stage progenitors did not impair Müller glia specification, but caused radial displacement of Müller glia nuclei into the outer retina and modest changes in glial gene expression. Loss of *Sox8/9* in mature Müller glia modestly increased proliferation post-injury without inducing neurogenic competence. These findings suggest that Sox8/9 are dispensable for gliogenesis and repression of neurogenic competence, but are essential for proper laminar positioning and maturation of retinal Müller glia.

## Introduction

Cell fate specification in the developing nervous system is largely directed by temporal patterning and involves a process known as developmental competence. This competence is characterized by dynamic changes in gene expression and regulation throughout developmental neurogenesis that mediate the ability of progenitors to generate individual cell types. Single-cell multiomic analysis of multiple regions in the developing vertebrate central nervous system (CNS) have identified gene regulatory networks that correlate directly with early and late stages of developmental competence (Lyu et al. 2021; Sagner et al. 2021; Kim et al. 2025; La Manno et al. 2021; Telley et al. 2019). These networks in turn include several transcription factors that show conserved patterns of temporal regulation across multiple CNS regions. Several of the transcription factors identified in the gene regulatory network active in late-stage progenitors were first implicated in controlling the generation of glia, which are invariably among the last-born cell types (Miller and Gauthier 2007; Frisén 2016). These include members of the nuclear factor I (NFI) and SoxE gene families (*Sox8/9/10*). Overexpression of both *Nfia* and *Sox9* induce astrogliogenesis in the brain, while loss of function compromises gliogenesis (Stolt et al., 2003) (Tchieu et al. 2019; Deneen et al. 2006; Takouda et al. 2021; Klum et al. 2018; Stolt et al. 2003) In the retina, overexpression of multiple NFI family members simultaneously suppresses generation of early-born cell types while promoting formation of late-born cell types and mitotic exit, while loss of function has the opposite effect (Lyu et al. 2021; Clark et al. 2019). Moreover, NFI factors and Sox9 frequently bind to common target sites controlling expression of genes that promote both late-stage temporal identity and gliogenesis (Kang et al. 2012). This has raised the possibility that NFI and SoxE factors may form an essential core component of the gene regulatory network that controls these processes (Zhang et al. 2023).

Since the major cell types of the retina are generated exclusively through dynamic changes in temporal patterning (Santos-França et al. 2023; Cepko 2014), it provides an ideal model to investigate this question. Müller glia (MG), the principal glial cells of the retina, emerge late in retinal development from retinal progenitor cells (RPCs) during a shift from neurogenesis to gliogenesis that coincides with increased expression of both NFI and SoxE factors (Lyu et al. 2021; Clark et al. 2019). The SoxE gene family includes *Sox8*, *Sox9*, and *Sox10*. While *Sox10* is not detectably expressed in neuroretina, *Sox8* and *Sox9* are robustly expressed in late-stage RPCs and their expression is maintained in mature MG (Clark et al. 2019; Lu et al. 2020) RPC-specific loss of function of *Sox9* has been reported to eliminate expression of mature MG markers without disrupting retinal integrity (Poché et al. 2008), while shRNA-mediated knockdown of either *Sox8* or *Sox9* in late-stage RPCs was reported to promote generation of rod photoreceptors at the expense of MG (Muto et al. 2009). This suggests that Sox8 and Sox9 may both be required to both promote retinal gliogenesis and suppress neurogenesis in late-stage RPCs.

This in turn raises the question as to whether sustained expression of *Sox8/9* in mature MG is required to repress neurogenic competence. Both loss of function of NFI factors and disruption of Notch signaling in mature MG induce neurogenic competence and lead to the generation of late-born inner retinal neurons, and in the case of Nfia/b/x, also induce limited MG proliferation (Le et al. 2024; Hoang et al. 2020). In both cases, downregulation of *Sox8/9* expression is also observed. This led us to hypothesize that these SoxE factors are themselves downstream targets of NFI and Notch signaling, and that their sustained expression may inhibit efficient MG reprogramming. We suspected that Sox8 and Sox9 may not only function as markers of mature MG identity, but may also act directly to enforce glial quiescence and suppress latent neurogenic programs.

In this study, we investigated the role of Sox8 and Sox9 in both developing and mature MG using both overexpression in early-stage RPCs and conditional loss of function in both late-stage RPCs and mature MG. While overexpression of *SOX8/9* either individually or in combination promoted formation of later-born photoreceptor precursors at the expense of early-born retinal ganglion cells as expected, we did not observe any changes in retinal cell fate following *Sox8/9* loss of function in late-stage RPCs either individually or in combination. Instead, we observed displacement of MG nuclei into the outer nuclear layer, along with a modest reduction in a subset of mature MG markers. Selective disruption of *Sox8/9* in mature MG likewise did not induce glial-derived neurogenesis, in either unstimulated or injury-induced condition, but did result in a modest increase in proliferation immediately following excitotoxic injury. Single-cell multiomic analysis of mutant cells revealed widespread changes in expression of genes controlling cell adhesion and migration in addition to mature glial markers, along with reduced accessibility of chromatin regions targeted by Sox8/9. We conclude that while Sox8/9 influence gene expression and nuclear positioning in MG, that they are ultimately dispensable for retinal gliogenesis.

## Results

### *Sox8* and *Sox9* are expressed in late-stage RPCs and mature Müller glia

We first sought to confirm previous reports of the cellular expression pattern of *Sox8* and *Sox9* in the developing retina. Analyzing scRNA-Seq data collected across the full time course of mouse retinal development (Fig. S1a), we find that both *Sox8* and *Sox9* show substantially higher expression in late-stage RPCs relative to early-stage RPCs, with this being particularly so for *Sox8*. Sustained expression is observed in differentiated MG at postnatal day (P)14, but not detected in retinal neurons at this age (Fig. S1b). Cellular expression levels of both *Sox8* and *Sox9* peak from E18-P2, in line with the transition from early- to late-stage identity, which occurs around E17 in mice (Lyu et al. 2021; Clark et al. 2019) (Fig. S1c). In adult retina, *Sox8* and *Sox9* expression is maintained in MG, with limited levels also observed in retinal astrocytes but not in subtypes of retinal neurons (Fig. S1d).

### Overexpression of *SOX8/9* in early-stage RPCs suppresses generation of early-born cell types

To determine whether overexpression of *Sox8/9* was sufficient to regulate temporal patterning in early-stage RPCs, we electroporated early-stage developing retinal explants with either control pCAGIG or plasmids overexpressing *SOX8* and *SOX9*, either individually or in combination (Fig. 1A). The explants were electroporated at embryonic day (E)14 and cultured until E19 (P0), with an EdU pulse delivered at E16 (Fig. 1B). We then used immunohistochemistry to analyze changes in cell composition and proliferation in GFP-positive electroporated cells. Overexpression of either *SOX8* or *SOX9* resulted in a significant reduction in the percentage of GFP-positive retinal ganglion cells (RGCs), which was not further increased by overexpression of both in combination (Fig. 1C, D). In contrast, we observed increased numbers of GFP and Otx2-positive photoreceptor precursor cells, with this effect being statistically significant when *SOX8* or both *SOX8*/*9* were overexpressed together (Fig. 1E, F). This matches the shift in temporal identity observed following overexpression of *Nfia/b/x* in early-stage RPCs (Lyu et al. 2021). We also observed a statistically significant decrease in the proportion of GFP-positive amacrine cells for each experimental condition compared to the control, although, interestingly, this effect was strongest with *SOX8* overexpression (Fig. 1 G, H). Fewer GFP and Sox2-positive RPCs were observed with *SOX8* and combined *SOX8*/*9* overexpression (Fig. S2a, b).

**Fig. 1:**
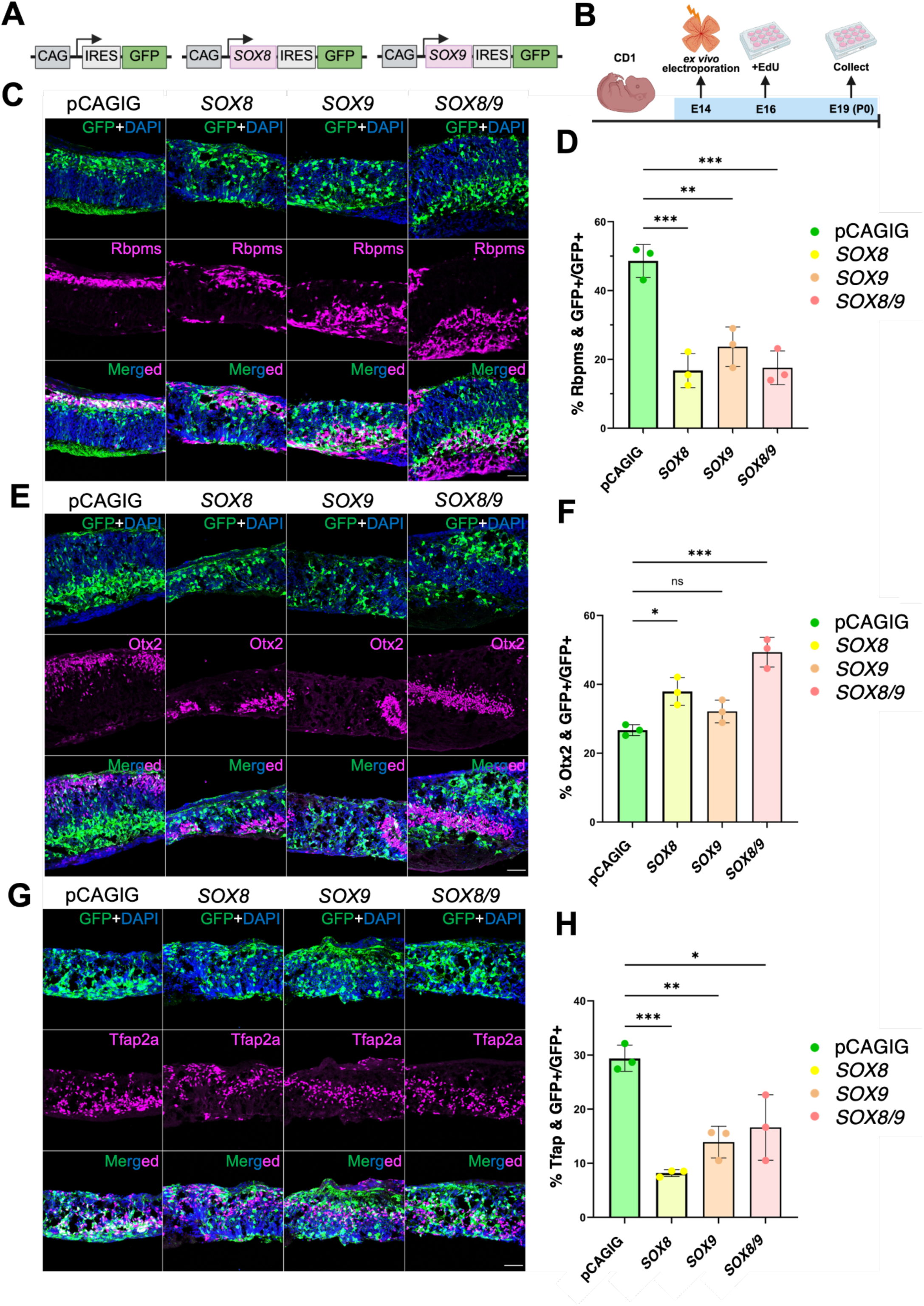
Overexpression of *SOX8* and *SOX9* in retinal explant cultures. **(A)** Schematic of the constructs used to induce expression of green fluorescent protein (*GFP)*, *SOX8*, and *SOX9*. **(B)** Schematic of the experimental workflow. P0 retinal explants were collected following *ex vivo* electroporation at E14 with control (pCAGIG), *SOX8*, *SOX9,* or pooled *SOX8* and *SOX9* constructs. **(C, E, G)** Representative immunostaining for GFP and retinal cell type markers **(C)** Rbpms (retinal ganglion cells), **(E)** Otx2 (photoreceptors), and **(G)** Tfap2a (amacrine cells) in electroporated retinal explants collected at P0. Scale bars, 50 μm. DAPI, 4′,6-diamidino-2-phenylindole. **(D, F, H)** Quantification of mean percentage ± SD of GFP-positive electroporated cells that co-label with **(D)** Rbpms, **(F)** Otx2 or **(H)** Tfap2a for each condition. Significance was determined via one-way ANOVA with Tukey’s test: \**P* < 0.0332, \*\**P* < 0.0021, \*\*\*\**P* < 0.0001. Each data point was calculated from an individual retina.

Increased GFP and Mki67-positive cells were observed for all experimental conditions, although this effect was only significant with *SOX8* overexpression (Fig.S2 c, d). In contrast, a statistically significant increase in EdU incorporation in GFP-positive cells was observed following overexpression of both *SOX8* and *SOX8*/*9* in combination, while increased EdU incorporation with *SOX9* overexpression did not reach statistical significance (Fig. S2e, f). Overall, these results may reflect transient stimulation of proliferation in neurogenic RPCs followed by cell cycle exit and depletion of primary RPCs, again consistent with a shift towards later-stage temporal identity.

### Loss of function of either *Sox8* or *Sox9* in late-stage RPCs leads to radial displacement of MG nuclei to the ONL

We first attempted to generate RPC-specific conditional mutants of *Sox8* and *Sox9* by crossing to the RPC-specific Chx10-Cre transgenic line (Rowan and Cepko 2004). We successfully generated *Chx10-Cre;Sox9^lox/lox^* mice and collected retina at P14 and P120 for scRNA-Seq analysis (Fig. S3a, b). Efficient deletion of exons 2 and 3 was observed in *Sox9*-deficient retinas (Fig. S3c), and all major retinal cell types, including MG, were identified at both timepoints (Fig. S3d, e). Direct comparison of gene expression profiles between control and *Sox9*-deficient MG revealed 497 differentially expressed genes (104 upregulated, 393 downregulated) at P14 and 682 differentially expressed genes (293 upregulated, 389 downregulated) at P120 (Table ST1). Small defects in Sox family (*Sox2*, *Sox5*, *Sox6*) and MG-specific (*Glul*, *Abca8a*) gene expression, along with slight induction of genes that promote neurogenesis (*Ascl1*, *Btg2*), were observed at P14, with an even more subtle effect observed at P120 (Fig. S3f-h). Gene set enrichment analysis of differentially expressed genes in P14 Sox9-deficient MG showed slight activation of relevant gene sets including neuron differentiation and generation of neurons (Fig. S3i). However, immunohistochemical analysis revealed no obvious defects in retinal neurogenesis, glial specification, or glial morphology (data not shown). We were also unable to generate any homozygous *Chx10-Cre;Sox8^lox/lox^* pups despite many rounds of breeding. We therefore instead selectively disrupted *Sox8* and *Sox9* in late-stage RPCs using both the *RaxCreER* knockin and the *GlastCreER* transgenic line, both of which have been previously shown to work effectively for this purpose (de Melo et al. 2012; Pak et al. 2014), in combination with the Cre-inducible *R26-CAG-lsl-Sun1-GFP* line, which labels the nuclear envelope (Mo et al. 2015). Cre recombination was induced in pups via intramuscular injection of tamoxifen from P3-5, and retinas were collected for immunohistochemical analysis at P60 (Fig. 2A-C). Both treatments efficiently induced loss of Sox8 immunoreactivity (Fig. 2D), and also led to mosaic loss of Sox9 immunoreactivity (Fig. 2E). Loss of function of either gene leads to displacement of MG nuclei into the outer nuclear layer as detected by Sox2 immunoreactivity, with stronger phenotype observed for *Sox8* mutants (Fig. 2F, G), although this may reflect the more efficient Cre-dependent loss of function of this gene. All displaced cells expressed Sox2, indicating that glial identity is preserved, in contrast to previous reports (Poché et al. 2008).

**Fig. 2.**
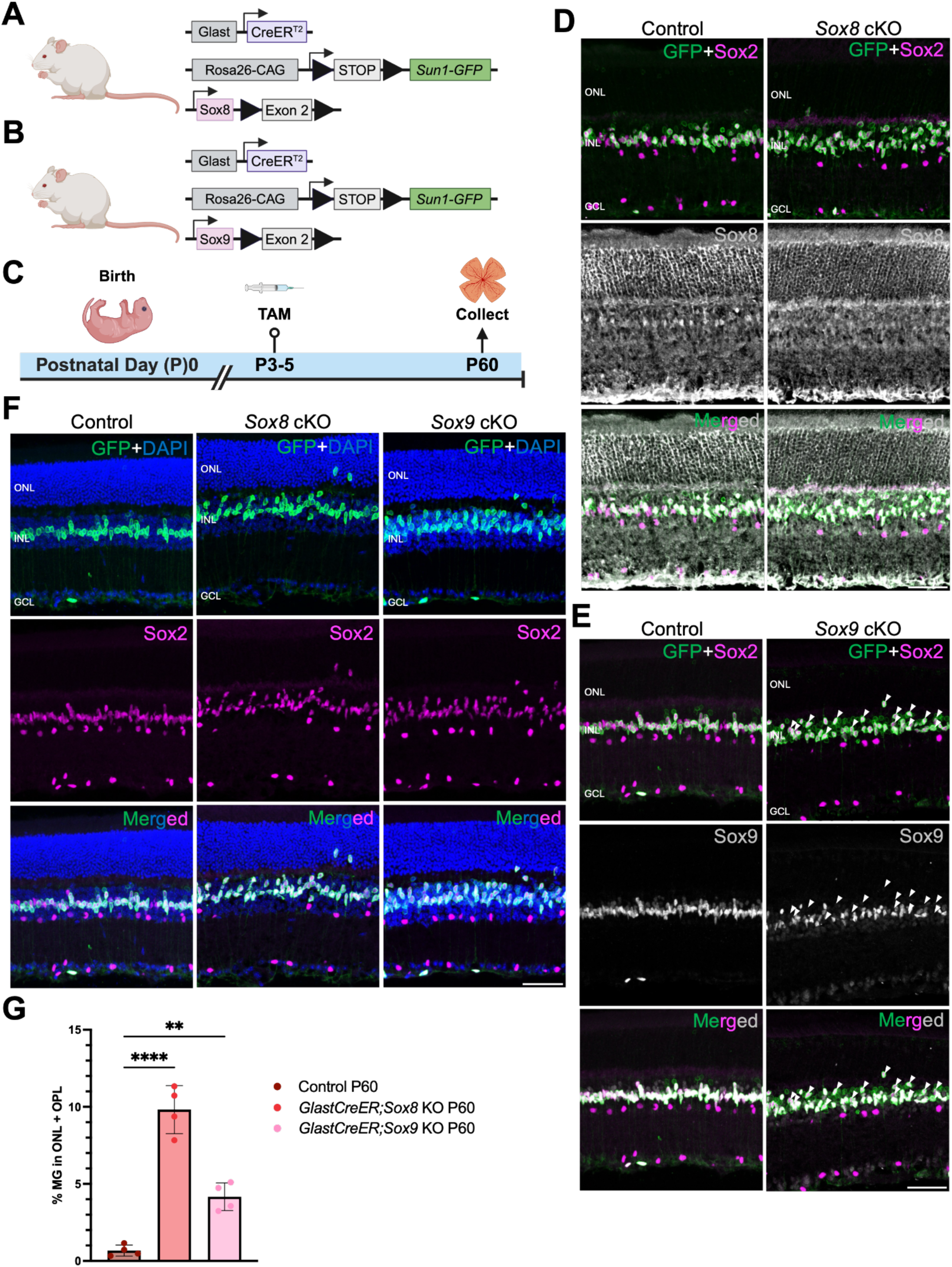
Selective loss of function of either *Sox8 or Sox9* in late-stage RPCs leads to radial displacement of MG nuclei to the ONL/OPL. (A and. **B)** Schematic of the transgenic constructs used to induce deletion of *Sox8* or *Sox9* specifically in late-stage RPCs/MG. Cre-mediated removal of Exon 2 of *Sox8* or *Sox9* leads to premature stop codon formation and disrupts the expression of the Sox8 or Sox9 protein. **(C)** Schematic of the experimental workflow. **(D)** Representative immunostaining for green fluorescent protein (GFP), Sox8, and Sox2 in control and *Sox8-*deleted retinas collected at postnatal day (P)60. **(E)** Representative immunostaining for GFP, Sox9, and Sox2 in control and *Sox9-*deleted retinas collected at P60. White arrowheads indicate co-labeled GFP and Sox2-positive cells that are Sox9-negative. **(F)** Representative immunostaining for GFP and Sox2 in control and either *Sox8* or *Sox9-*deleted retinas collected at P60. ONL, outer nuclear layer; INL, inner nuclear layer; GCL, ganglion cell layer. Scale bars, 50 μm. DAPI, 4′,6-diamidino-2-phenylindole. **(G)** Quantification of mean percentage ± SD of GFP-positive MG displaced to either the outer nuclear layer or outer plexiform layer. Significance was determined via two-way ANOVA with Tukey’s test: \**P* < 0.0332, \*\**P* < 0.0021, \*\*\*\**P* < 0.0001. Each data point was calculated from an individual retina.

### Combined loss of function of both *Sox8* and *Sox9* in late-stage RPCs leads to increased numbers of radially displaced MG

We next tested the effects of combined loss of function of *Sox8* and *Sox9* in late-stage RPCs, again using both *RaxCreER* and *GlastCreER* and the *R26-CAG-lsl-Sun1GFP* line (Fig. 3A, E) and administering tamoxifen from P3-5. The retinas were then collected at P16 and P60 or P90 for immunohistochemical analysis (Fig. 3B, F). In both cases, we observed substantial levels of MG displacement at P16, with 15% of GFP-positive MG nuclei radially displaced in *RaxCreER*;*Sox8^lox/lox^;Sox9^lox/lox^*;*R26-CAG-lsl-Sun1GFP* (Fig. 3C, D) and 25% of *GlastCreER*;*Sox8^lox/lox^;Sox9^lox/lox^* ;*R26-CAG-lsl-Sun1GFP* (Fig. 3G-I) displaced into the ONL. At P90, this increased to 31% in *RaxCreER*;*Sox8^lox/lox^;Sox9^lox/lox^*;*R26-CAG-lsl-Sun1GFP* mice (Fig. 3D), but no significant age-dependent increase in displacement is seen with *GlastCreER*;*Sox8^lox/lox^;Sox9^lox/lox^* ;*R26-CAG-lsl-Sun1GFP* mice at P60 (Fig. 3I).

**Fig. 3.**
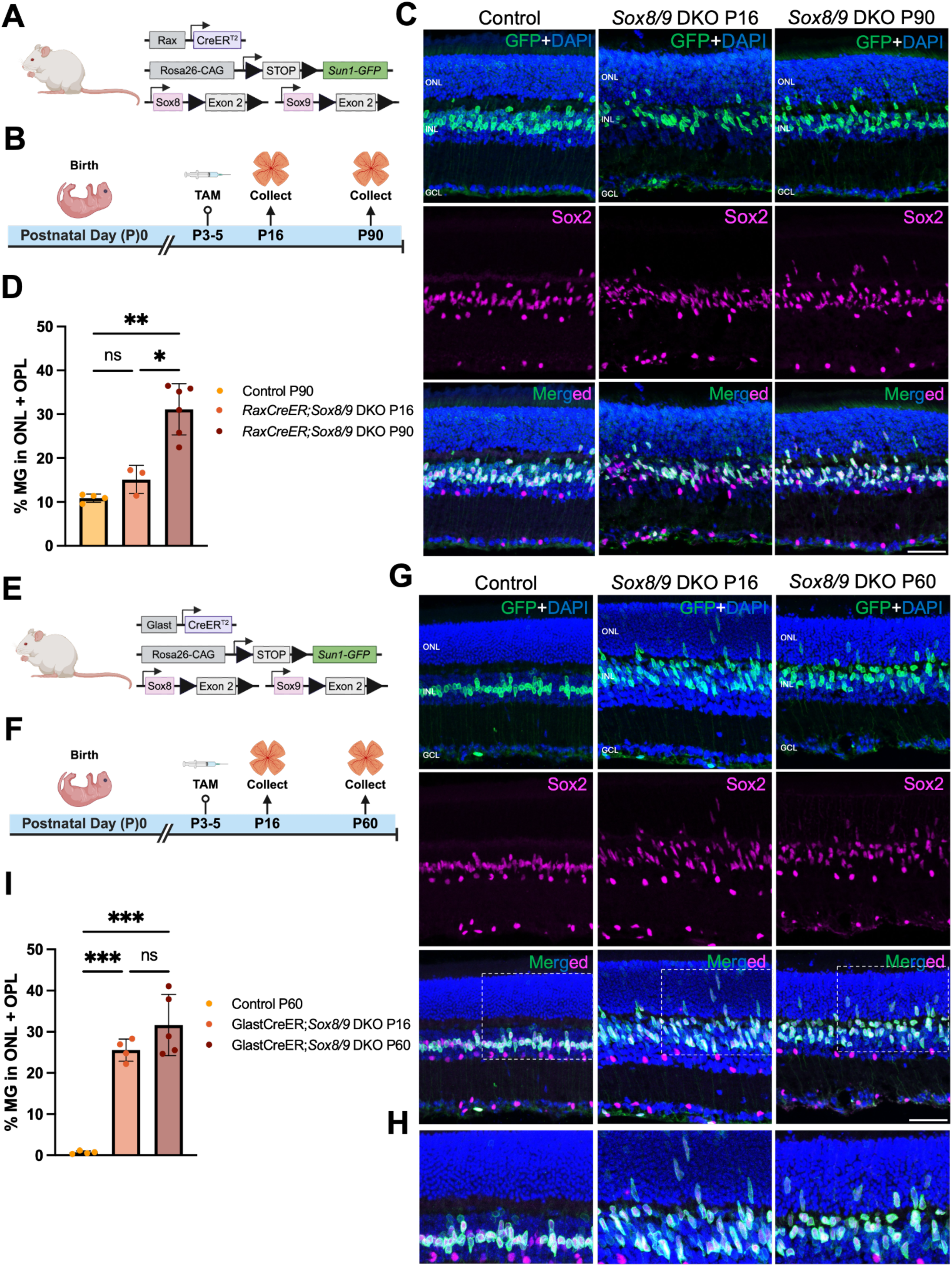
Selective loss of function of *Sox8/9* during late-stage retinal development enhances MG displacement to the ONL/OPL. (A and. **E)** Schematic of the transgenic constructs used to induce deletion of *Sox8/9* specifically in late-stage RPCs/MG. Cre-mediated removal of Exon 2 of *Sox8/9* leads to premature stop codon formation and disrupts the expression of the Sox8/9 proteins. **(B and F)** Schematic of the experimental workflow. **(C, G-H)** Representative immunostaining for GFP and Sox2 in control and *Sox8/Sox9*-deleted retinas collected at P16 and P90 **(B)** or P60 **(G,H)**. Scale bars, 50 μm. **(D and I)** Quantification of mean percentage ± SD of GFP-positive MG displaced to either the ONL or OPL. Significance was determined via two-way ANOVA with Tukey’s test: \**P* < 0.0332, \*\**P* < 0.0021, \*\*\**P* < 0.0002. Each data point was calculated from an individual retina. Scale bars, 50 μm.

At all timepoints, we also observe that most displaced MG nuclei show elongated morphology, in contrast to GFP-positive cells that remained in the INL (Fig. 3C, G-H). We unexpectedly observed that 10% of MG in *RaxCreER*;*R26-CAG-lsl-Sun1GFP* control mice also show nuclear displacement at P90 (Fig. 3C, D). Since the *RaxCreER* knock-in line disrupts expression at the endogenous *Rax* locus, this suggests that this may represent a previously unreported effect of *Rax* heterozygosity, which has been observed in adult hypothalamus (Miranda-Angulo, et al. 2013). For this reason, along with the more efficient overall disruption of *Sox9* expression, we therefore used the *GlastCreER*;*Sox8^lox/lox^;Sox9^lox/lox^* for the remainder of these studies.

### Combined loss of function of both *Sox8* and *Sox9* in late-stage RPCs does not grossly compromise glial-specific gene expression or retinal lamination

We next used immunostaining to determine whether there were any obvious disruptions in MG-specific gene expression or morphology in P60 *GlastCreER*;*Sox8^lox/lox^;Sox9^lox/lox^* ;*R26-CAG-lsl-Sun1GFP* mice. We observed a reduction in glutamine synthetase staining in the inner nuclear layer, although signal was preserved in both apical and basal MG processes (Fig. S4a). In contrast, we did not observe any obvious changes in Aqp4 distribution (Fig. S4b), nor did we observe any induction of GFAP (Fig. S4c) or loss of Hes1 expression (Fig. S4d). Radial displacement of MG nuclei did not affect expression of markers of the inner limiting membrane such as Lama1 (Fig. S4e) or ZO-1 (Fig. S4f). Finally, in contrast to *Nfia/b/x* or *Rbpj*-deficient MG (Le et al. 2024; Hoang et al. 2020), we do not observe any displaced cells in the ONL expressing either the inner retinal neuronal marker Pax6 (Fig. S4g) or the bipolar cell and photoreceptor marker Otx2 (Fig. S4h).

### Single-cell multiome and CUN&RUN analysis identify Sox8/9-regulated genes

To comprehensively analyze changes in gene expression and chromatin accessibility resulting from loss of function of *Sox8/9*, we performed combined snRNA- and ATAC-seq multiome analysis on GFP-positive cells that were FACS-isolated from P17 *GlastCreER*;*Sox8^lox/lox^;Sox9^lox/lox^* ;*R26-CAG-lsl-Sun1GFP* mice, in parallel with age-matched *GlastCreER*;*R26-CAG-lsl-Sun1GFP* controls (Fig. 4A). While GFP labeled both late-born rod photoreceptors and bipolar cells, as well as MG, we focused exclusively on MG for the purpose of this analysis. UMAP analysis of MG from control and *Sox8/9* mutants revealed extensive overlap in gene expression profiles, but substantial divergence of a subset of mutant MG (Fig. 4B).

**Fig. 4.**
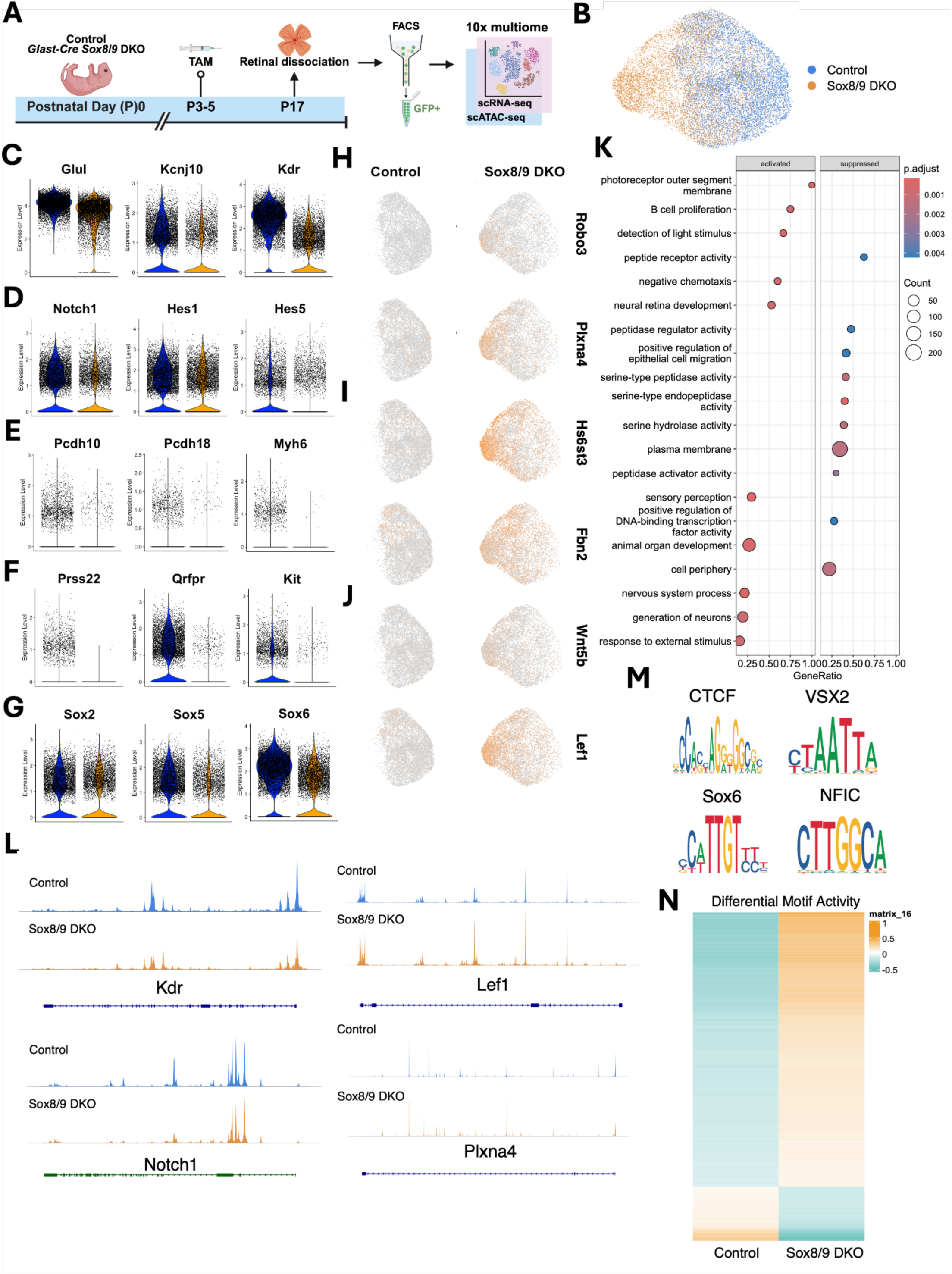
Multiome analysis of control and *Sox8/9* DKO Müller glia. **(A)** Schematic of the multiome experimental pipeline. **(B)** UMAP plot of multiome datasets showing the clustering of control and *Sox8/9*-deficient MG. **(C-G)** Violins plots showing decreased expression of **(C)** MG markers (*Glul*, *Kcnj10, Kdr*), **(D)** Notch signaling genes (*Notch1, Hes1, Hes5*), **(E)** regulators of cell adhesion (*Pcdh10*, *Pcdh18)* and myosin-dependent motility (*Myh6*), **(F)** cell migration (*Prss22*, *Qrfpr, Kit*), and **(G)** *Sox* genes (*Sox2*, *Sox5*, *Sox6*). **(H-J)** Feature plots showing increased expression of **(H)** regulators of cell migration and/or synaptogenesis (*Robo3*, *Plxna4*), **(I)** extracellular matrix genes (Hs6st3, *Fbn2*), and **(J)** Wnt signaling genes (*Wnt5b*, *Lef1*). **(K)** Gene set enrichment analysis of significant differentially expressed genes. **(L)** Coverage plots for a subset of differentially expressed genes from C and D. **(M)** Depleted motifs in *Sox8/9*-deficient MG. **(N)** Heatmap showing differential motif activity between conditions.

Direct comparison of gene expression profiles between control and *Sox8/9* mutant MG identified 2,336 differentially expressed genes, with 460 upregulated and 1,876 downregulated (Table ST2). We observe reduced expression of multiple MG-specific genes in the mutants, with significant reductions of *Glul* (Glutamine Synthetase), the potassium channel *Kcnj10*, and the VEGF receptor *Kdr* (Fig. 4C). Notch pathway genes (*Notch1*, *Hes1*, and *Hes5*) were also moderately downregulated (Fig. 4D). The most strongly downregulated genes were enriched for regulators of cell adhesion (*Pcdh10*, *Pcdh18*, *Alcam1*), cell migration (*Prss22*, *Kit*, *Qrfpr*), or myosin-dependent motility (*Myh6*, *Myof*) (Fig. 4E, F). Several other Sox family members, including *Sox5* and *Sox6*, were also strongly downregulated, with *Sox2* also showing a modest reduction (Fig. 4G).

Strongly upregulated genes included regulators of cell migration and/or synaptogenesis (*Robo3*, *Plxna4*), extracellular matrix (Hs6st3, *Fbn2*), or Wnt signaling (*Lef1*, *Wnt5b*) (Fig. 4H-J). Gene set enrichment analysis of differentially expressed genes identified several functional categories that matched these observations, including negative chemotaxis in upregulated genes and positive regulation of epithelial cell migration in downregulated genes (Fig. 4K). These are consistent with Sox8/9 playing a general role in regulating adhesion and radial positioning of Muller glia.

We next analyzed the scATAC-Seq data to identify differentially accessible regions associated with these genes. 3,924 differentially accessible regions were identified, with 1,109 showing increased accessibility and 2,815 having reduced accessibility (Table ST2). We observed a widespread and consistent reduction in accessibility in regulatory regions associated with downregulated genes (Fig. 4L). SOX family motifs, including Sox8 and Sox9 consensus sites, were the most strongly enriched motifs in regions that showed reduced accessibility in mutants, as expected (Fig. 4M, N). RORB and FOS/JUN motifs were also enriched in these regions. Motifs showing increased accessibility were enriched for CTCF and multiple Lhx family members, which likely correspond to Lhx2, which promotes gliogenesis and represses reactive gliosis (Zibetti et al. 2019; de Melo, Zibetti, et al. 2016; de Melo et al. 2012).

To determine whether any major changes in MG-specific gene expression were observed in older animals, we also performed scRNA-Seq analysis of P90 *RaxCreER*;*Sox8^lox/lox^;Sox9^lox/lox^* ;*R26-CAG-lsl-Sun1GFP* and *RaxCreER*;*R26-CAG-lsl-Sun1GFP* littermate controls (Fig. S5a). All major retinal cell types, including MG, were identified in similar proportions at this timepoint (Fig. S5b-d). Direct comparison of MG-specific gene expression profiles revealed some similar patterns of differential gene expression. This includes modest downregulation of several mature MG markers *(Glul, Kcnj10)*, Notch pathway components (*Notch1*, *Hes1*), cell migration genes (*Qrfpr*), and myosin-dependent motility genes (*Myof*), along with upregulation of genes associated with cell migration and/or synaptogenesis (*Plxna4*) and Wnt signaling (*Lef1*) (Fig.S5e). Sox family factors (*Sox2, Sox3, Sox4, Sox5*, *Sox6, Sox8, Sox9, Sox11*) were also generally moderately downregulated (Fig. S5f). 1,675 differentially expressed genes were identified, with 1,094 upregulated and 581 downregulated (Table ST3, Fig. S5g). However, gene set enrichment analysis of differentially expressed genes did not identify functional categories that matched these observations, which may correlate with the milder phenotype that we observed using the *RaxCreER* transgenic line (Fig. S5h).

We next performed CUT&RUN analysis to identify direct targets of Sox8 and Sox9 at both P2 and P17 in wildtype mouse retinas (Fig. 5A, Fig. S6A-E). The data was integrated with previously published scATAC-Seq data from P2 retina (Lyu et al. 2021) and P17 scATAC-Seq data from this study. We detected 5,616 Sox8/9 peaks in P2 retinas, with 437 of these peaks overlapping with differentially accessible regions (DARs) detected in P2 RPCs (Fig. 5B). Gene set enrichment analysis of these overlapping peaks revealed enrichment of multiple developmental processes, including regulation of neurogenesis, eye development, axon guidance, and cell fate commitment (Fig 5C). At both ages, we observed that both Sox8 and Sox9 mainly bind to DARs located within 1 kb of transcription start sites (TSS) (Fig. 5D,E). The great majority of Sox8/9 peaks associated with DARs present in P2 RPCs overlapped with H3K27ac, and in P17 MG with both H3K27ac and H3K4me3, indicating that they represent active promoter or enhancer elements (Fig. 5F,G). Loss of function of *Sox8/9* led to a dramatic reduction in accessible chromatin at these sites, indicating that Sox8/9 are essential for maintaining chromatin accessibility at their target sites (Fig. 5F).

**Fig. 5.**
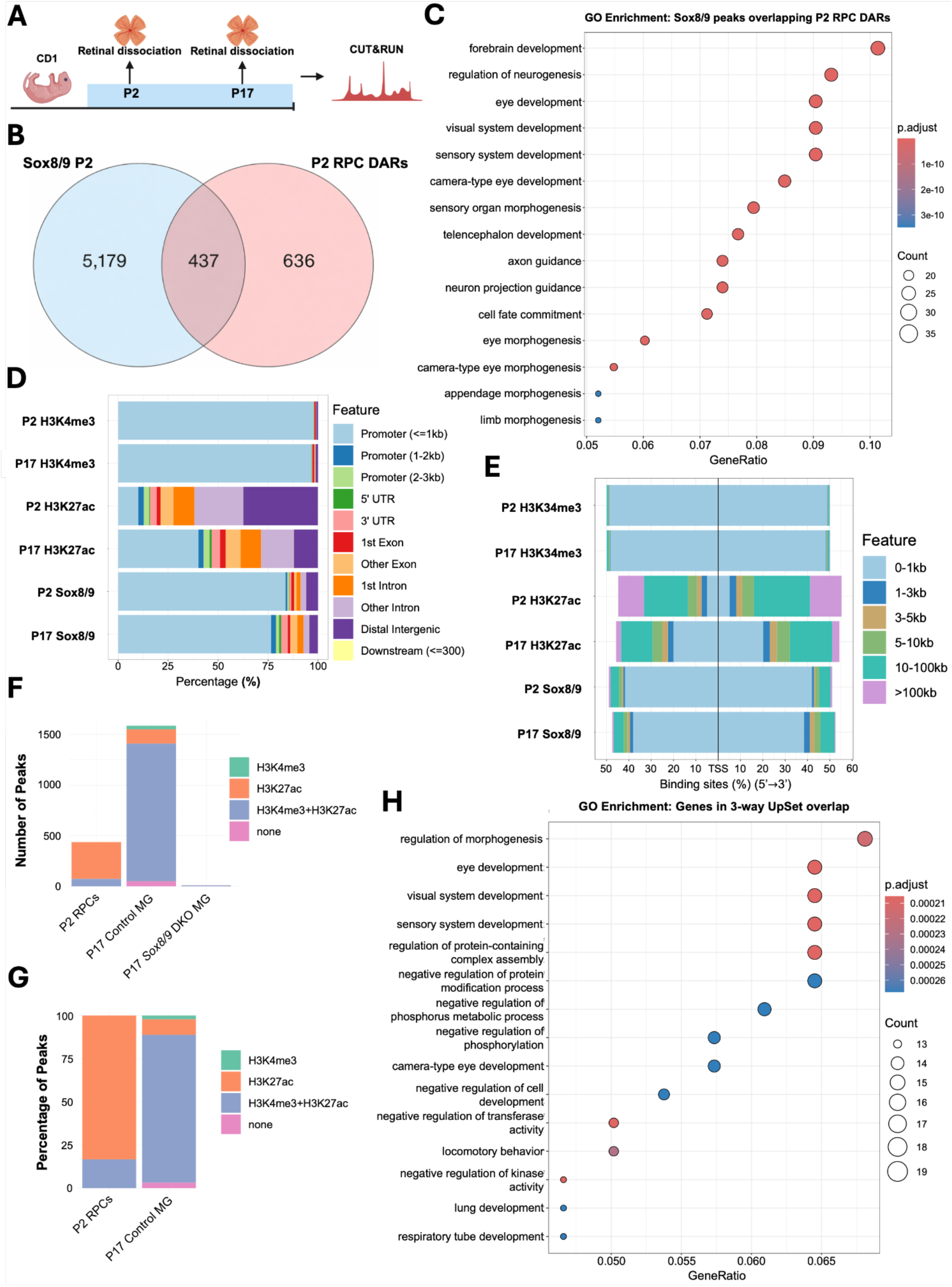
CUT&RUN analysis of Sox8/9 in P2 and P17 mouse retina. **(A)** Schematic of the CUT&RUN experimental pipeline. **(B)** Overlap of P2 Sox8/9 peaks and P2 RPC DARs. **(C)** GO enrichment of P2 Sox8/9 peaks and P2 RPC DARs. **(D)** Distribution of Sox8/9 binding loci in distinct gene regions at P2 and P17 compared to H3K27ac and H3K4me3 controls**. (E)** Distribution of Sox8/9 binding loci relative to transcription start site (TSS) at P2 and P17 compared to H3K27ac and H3K4me3 controls**. (F)** Number of Sox8/9 CUT&RUN peaks that overlap with enriched/differentially accessible regions in either P2 RPCs, P17 Control MGs, or P17 Sox8/9 cKO MGs. **(G)** Distribution of Sox8/9 CUT&RUN peaks that overlap with enriched/differentially accessible regions in either P2 RPCs or P17 Control MGs. **(H)** Gene ontology enrichment of 3-way overlapping *Sox8/9* DKO differentially expressed genes, differentially accessible regions, and P17 Sox8/9 peaks.

Gene set enrichment analysis of 3-way overlapping peaks showed enrichment of cellular processes relevant to the phenotype observed in *Sox8/9*-deficient MG, including regulation of morphogenesis, eye development, negative regulation of cell development, and locomotory behavior (Fig. 5H). We also saw that both Sox8 and Sox9 are preferentially associated with accessible chromatin regions that are enriched for both Sox family motifs and the RPC and MG-enriched homeodomain factor Lhx2, which both bulk and single-cell ATAC-Seq analysis (Lyu et al. 2021; Zibetti et al. 2019) (Table ST4). Motifs for KLF family transcription factors, multiple members of which are enriched in late-stage RPCs (Clark et al. 2019), are also observed, particularly at P2. Sox8/9 peaks overlap with motifs for multiple factors – including CTCF, RORB, and FOS/JUN – that also show reduced accessibility in scATAC-Seq analysis of P17 *Sox8/9* mutant MG (Fig. 4M,N). We observe that *Sox5/8/9* are all directly targeted by Sox8/9 at P2, and Sox8/9 binding sites are also enriched for other classes of transcription factors, notably Sp1 and ZBTB family members, whose expression is differentially regulated in *Sox8/9* mutants. Furthermore, at P2 and P17 ages, we see that Sox8 and/or Sox9 target sites directly target regulatory sequences associated with Notch pathway genes (*Notch1*, *Hes1/5*), as well as MG-specific genes (*Kcnj10*, *Kdr, Fxyd3*) and genes regulating cell migration and adhesion (Cdh2, Ddr1, *Pcdh10*, *Kit*) that are differentially expressed in *Sox8/9* mutants.

### Loss of function of *Sox8/9* in mature MG induces limited levels of proliferation but does not induce neurogenic competence

Finally, to investigate whether loss of function of *Sox8* and *Sox9* either individually or combination was sufficient to induce either proliferation or neurogenesis in MG, we administered tamoxifen-supplemented chow to P21 mice for three weeks (Fig. 6A). To create injury conditions that are permissive for glial-derived neurogenesis (Le et al. 2024; Jorstad et al. 2017), at one week following resumption of normal chow, we induced excitotoxic damage by intravitreal injection as described (Hoang et al. 2020) and pulsed with EdU for three consecutive days immediately following this. We then harvested retinas at 4 weeks post-injury to investigate injury-induced proliferation of GFP-positive MG. Similar to our model of *Sox8/9* loss of function in late-stage RPCs, we observed efficient *Sox8* deletion following the TAM diet regimen, while *Sox9* deletion was mosaic (Fig. 6B). While we observe minimal levels of EdU incorporation in control animals, we observe a significant increase in EdU induction in both *Sox8* and *Sox8/9* double mutants (Fig. 6C-D). Both EdU-positive and EdU-negative *Sox8/9*-deficient GFP-positive cells are immunoreactive for Sox2 (Fig. 6E). No GFP-positive cells express Ki67, indicating that EdU incorporation occurred transiently immediately following injury (Fig. 6F). In addition, we observe that GFP-positive MG also largely continue to express the Notch pathway gene Hes1 (Fig. S7a), do not express the neurogenic bHLH factor Ascl1 (Fig. S7b), and do not show any co-expression of GFP with selective markers of late-born retinal neurons (Fig. S7c,d), indicating that *Sox8/9*-deficient MG lack neurogenic competence even following excitotoxic injury.

**Fig. 6.**
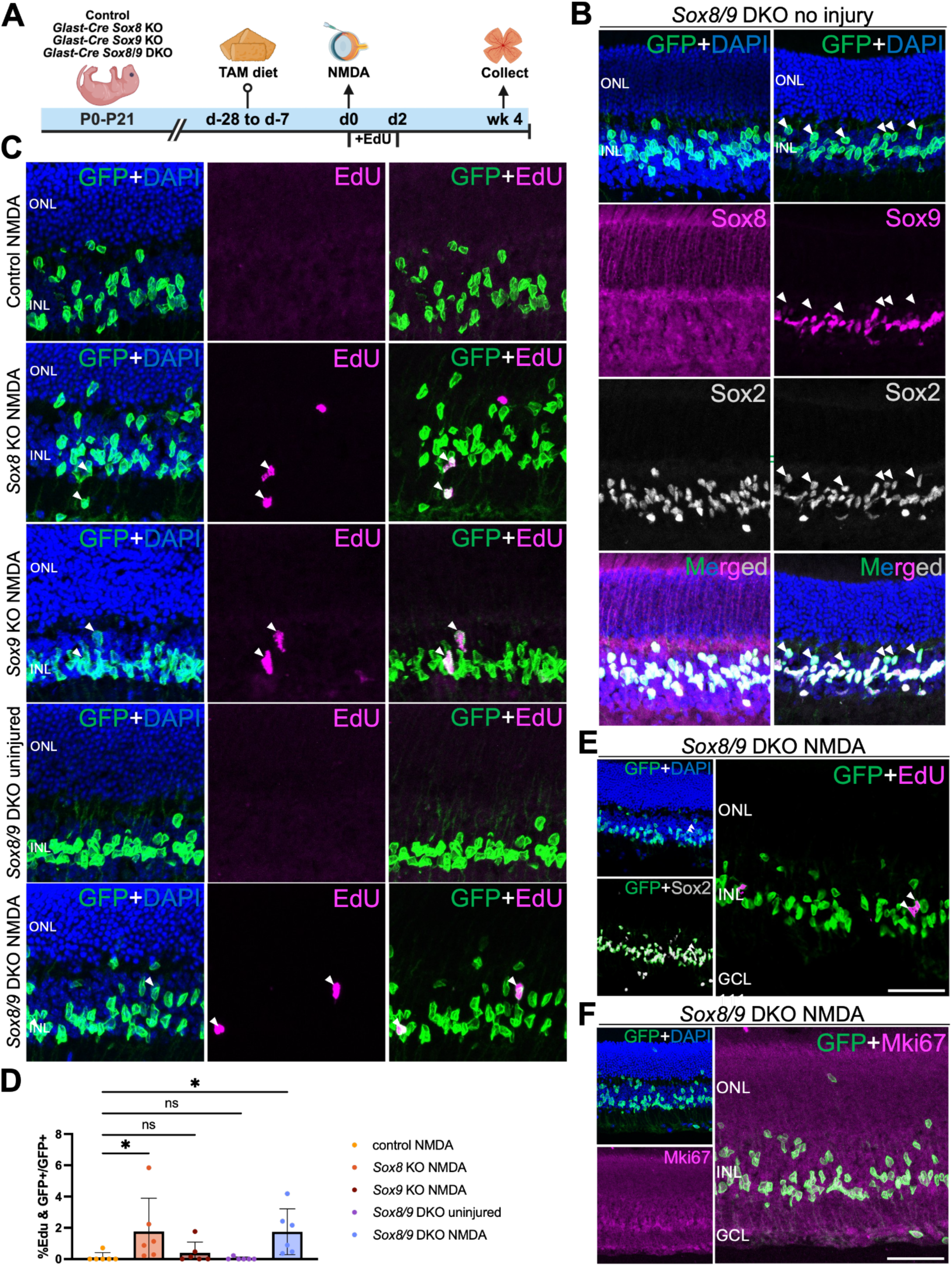
Selective loss of function of *Sox8/9* in mature MG stimulates proliferation with retinal injury. **(A)** Schematic of the experimental workflow. **(B)** Representative immunostaining for GFP and Sox8 or Sox9 in uninjured *Sox8/9*-deleted retinas five weeks after finishing the TAM diet. White arrowheads indicate co-labeled GFP and Sox2-positive cells that are Sox9-negative. **(C)** Representative immunostaining for GFP and EdU in control, *Sox8*, *Sox9*, and *Sox8/9*-deleted retinas collected four weeks following retinal injury. White arrowheads indicate co-labeled GFP-positive and EdU-positive cells. **(D)** Quantification of mean percentage ± SD of EdU-positive MG. Significance was determined via two-way ANOVA with Tukey’s test: \**P* < 0.0332. Each data point was calculated from an individual retina. **(E and F)** Representative immunostaining for GFP and Sox2 **(E)** and Mki67 **(F)** in *Sox8/9*-deleted retinas collected four weeks following retinal injury. Scale bars, 50 μm.

## Discussion

Transcription factors that control late-stage temporal identity in neural progenitors play a central role in the specification of late-born neurons and glia, and in repressing proliferative and neurogenic competence in mature glia (Clark et al. 2019). In light of both their expression pattern and their predicted function in gene regulatory networks reconstructed from single-cell multiomic analysis (Lyu et al. 2021), SoxE factors are prime candidates for playing a central role in this process. In this study, we systematically reassess the *in vivo* roles of the SoxE factors Sox8 and Sox9 across retinal development and in mature MG using temporally controlled genetic perturbations and single-cell genomic approaches. We find that overexpression of *SOX8* and/or *SOX9* in early retinal progenitors is sufficient to bias temporal output toward later-born fates, consistent with their predicted role in promoting late-stage temporal identity. Genetic loss of function of *Sox8* and *Sox9*, either individually or in combination, does not alter retinal cell fate specification, refining prior models that proposed an essential role for SoxE factors in MG specification (Muto et al. 2009; Poché et al. 2008). Likewise, we do not observe induction of neurogenesis in *Sox8/9*-deficient MG, even under injury paradigms previously shown to robustly induce neurogenesis following loss of function of *Nfia/b/x* or Notch signaling (Le et al. 2024; Hoang et al. 2020). Instead, we observe a reproducible but restricted phenotype characterized by radial displacement of a subset of MG, modest changes in mature glial gene expression, and a limited injury-induced proliferative response that does not progress to neurogenesis.

Our observation that *SOX8/9* overexpression suppresses generation of early-born cell types is consistent with these genes being able to modulate temporal patterning at this age, but does not imply that they can also do so when strong determinants of temporal identity such as *Nfia/b/x* are expressed. Indeed, previous studies have failed to observe any increase in retinal gliogenesis following postnatal overexpression of either *Sox8* or *Sox9* (de Melo, Clark, et al. 2016). Our analysis of *Sox8/9* mutants are also seemingly in contradiction with previous reports of an essential role for Sox8 and Sox9 in promoting retinal gliogenesis (Muto et al. 2009; Poché et al. 2008). Using multiple temporally and cell-type–restricted genetic strategies, we do not observe the loss of MG previously reported following *Sox9* disruption, suggesting that earlier findings may reflect differences in timing or targeting strategies. Another study that used shRNA to reduce *Sox8* and *Sox9* expression reported an increased photoreceptor generation and a corresponding reduction in MG generation (Muto et al. 2009). This, however, was conducted from E17.5 onwards, in contrast to our own work in which *Sox8/9* were disrupted at later stages and this, in addition to potential off-target effects of the constructs used, may reconcile our observations. Finally, a more recent study reported rapid rod photoreceptor degeneration following global conditional disruption of *Sox9* in adult mice (Hurtado et al. 2024). This in turn may reflect a disruption of the visual cycle due to loss of *Sox9* function in the retinal pigment epithelium, where it plays an essential role in regulating expression of these genes (Masuda et al. 2014).

The most prominent effect of *Sox8/9* disruption that we observed was the radial displacement of the nuclei of a subset of MG into the outer nuclear layer, along with the adoption of an elongated nuclear morphology in these displaced cells. While Sox9 promotes migration of non-CNS cell types – most notably tumor cells (Yang et al. 2023; Wang et al. 2021; Rockich et al. 2013; Larsimont et al. 2015) – neither it nor Sox8 have been linked to control of lamination or radial migration in the CNS. This is reminiscent of the phenotype observed following postnatal deletion of *Sox2* (Bachleda et al. 2016), although the severity is substantially greater following *Sox8/9* deletion. ScRNA-Seq analysis shows altered expression of genes encoding cell adhesion, extracellular matrix formation, cell migration, and myosin-based motility. Loss of function of Sox8/9 target genes such as *Kit*, *Dpysl3*, and *Rnd2* have been linked to defects in cell migration in the developing CNS (Guijarro et al. 2013; Tanaka et al. 2012; Nakamura et al. 2006), and loss of function of cell adhesion molecules such as *Pchd10* and *Pcdh18* regulate both cell adhesion and migration (Zhen et al. 2023; Aamar and Dawid 2008). These data support a model in which Sox8/9 function within a distinct gene regulatory module that, in combination with Sox2 and Sox5/6, stabilizes MG morphology and nuclear positioning, rather than specifying fate or neurogenic competence. Future work dissecting how Sox8/9 interact with other components of late-stage glial regulatory networks will further clarify how these transcriptional regulatory networks stabilize glial identity and morphology.

Gene regulatory networks controlling temporal patterning and glial maturation are highly redundant, and our findings indicate that Sox8/9 act in a subsidiary or dependent capacity relative to core regulators such as NFI factors and Notch signaling. In contrast to perturbations of NFI factors or Notch signaling, which are both necessary and sufficient to induce robust MG reprogramming (Clark et al. 2019; Lyu et al. 2021; Le et al. 2024; Hoang et al. 2020), loss of *Sox8/9* produces a qualitatively distinct and more constrained phenotype.

This implies that Sox8/9 have a subsidiary and dependent function in controlling both processes, which is supported by single-cell RNA-Seq data showing reduced expression of both genes in *Nfia/b/x* and *Rbpj*-deficient MG (Le et al. 2025; Hoang et al. 2020). Sox8/9 in turn appear to control expression of other transcription factors that are part of this late-stage RPC and glial-specific network, such as *Sox5/6*, which in turn inhibit proliferation and modulate gliogenesis in other CNS regions (Martinez-Morales et al. 2010; Stolt et al. 2006; Lefebvre 2010; Scheel et al. 2005). A more complete understanding of the hierarchical organization of the transcription factor networks that maintain glial quiescence and identity will help guide ongoing efforts aimed at efficiently and safely inducing glial reprogramming for treatment of neurodegenerative and blinding diseases (Blackshaw and Cayouette 2025; Langhe and Pearson 2020; Goldman 2014).

## Acknowledgements

### Data availability

All bulk RNA-Seq and scRNA-Seq data are available in GEO as GSE312862 and GSE311551. The code used in this paper is available at https://github.com/csanti88/sox8sox9_dko_pannullo_2025.

## Acknowledgements

This work was supported by NIH grant R01EY036173 to S.B.

---

**Figure S1:**
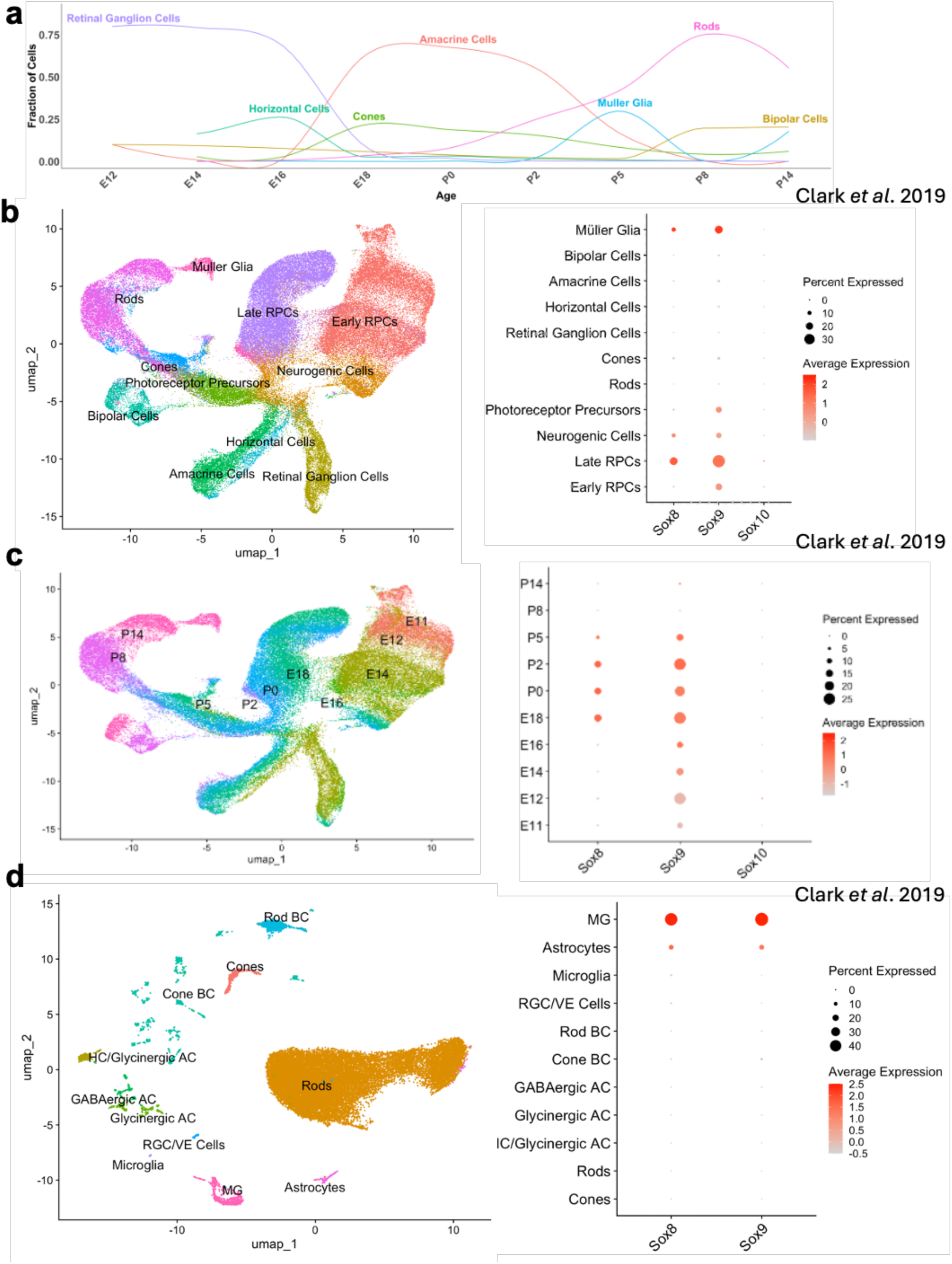
**Expression of SoxE family transcription factors in the developing and mature mouse retina**. **(a)** Line graph showing the birth of major retinal cell types over the course of retinal development. **(b and c)** UMAP plot of scRNA-seq data from developing mouse retina and dot plot showing cellular expression levels of SoxE family transcription factors in each major cell type **(b)** and time point **(c)**. **(d)** UMAP plot of scRNA-seq data from adult mouse retina and dot plot showing cellular expression levels of *Sox8* and *Sox9* in each major cell type.

**Fig. S2:**
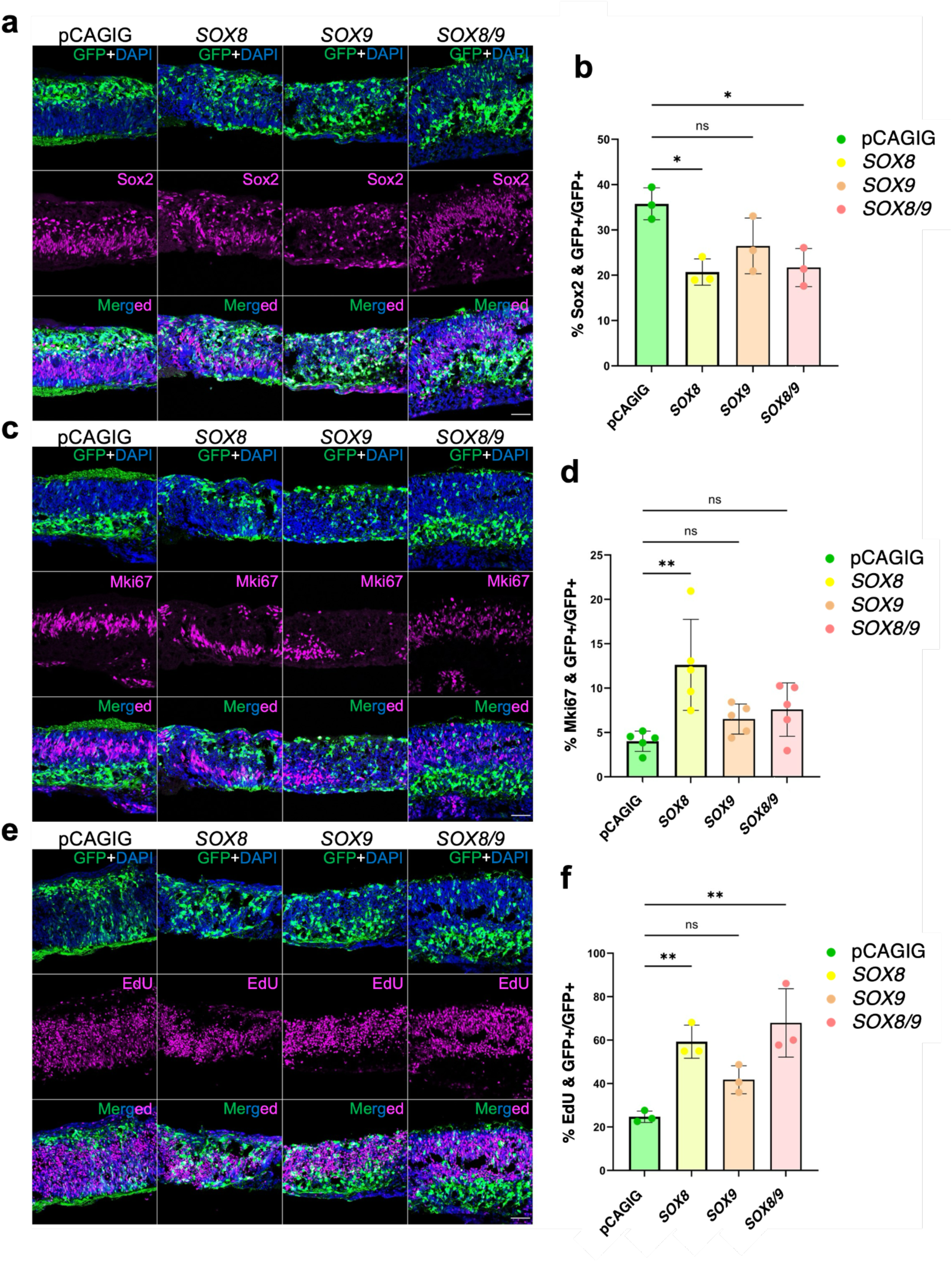
Overexpression of *SOX8* and *SOX9* in retinal explant cultures. (a, c, **e)** Representative immunostaining for GFP and **(a)** Sox2 (retinal progenitor cells and amacrine cells), **(c)** Mki67 (actively proliferating cells), and **(e)** EdU pulsed at E16 and chased until explant collection at P0. Scale bars, 50 μm. DAPI, 4′,6-diamidino-2-phenylindole. **(b, d, f)** Quantification of mean percentage ± SD of GFP-positive electroporated cells that co-label with **(b)** Sox2, **(d)** Mki67 or **(f)** EdU for each condition. Significance was determined via one-way ANOVA with Tukey’s test: \**P* < 0.0332, \*\**P* < 0.0021, \*\*\*\**P* < 0.0001. Each data point was calculated from an individual retina.

**Figure S3:**
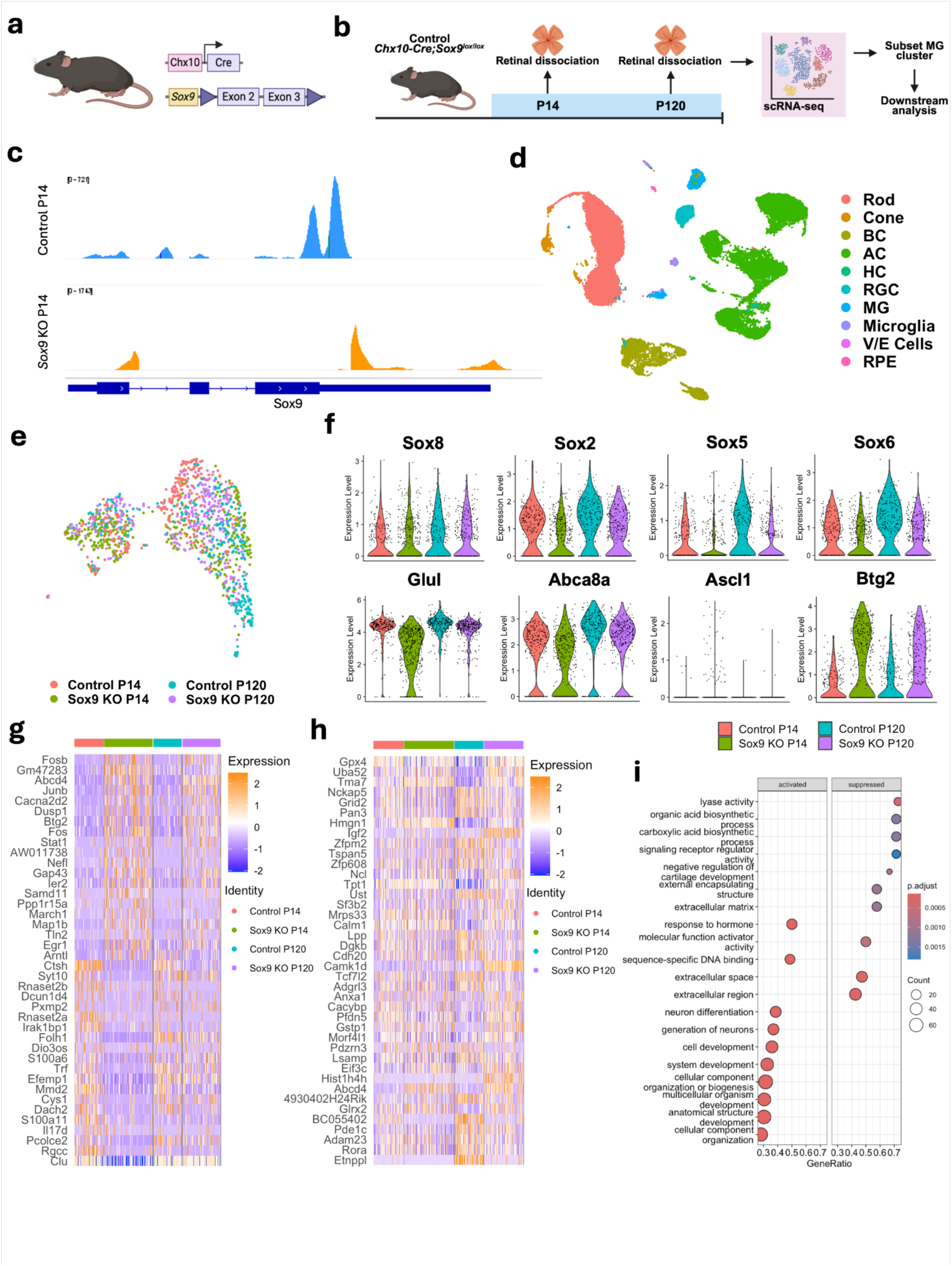
scRNA-seq analysis of control and *Chx10;Sox9^lox/lox^* Müller glia. **(a)** Schematic of the transgenic constructs used to induce deletion of *Sox9* specifically in RPCs. Cre-mediated removal of Exons 2 and 3 of *Sox9* leads to premature stop codon formation and disrupts the expression of the Sox9 protein. **(b)** Schematic of the scRNA-seq experimental pipeline. **(c)** scRNA-seq read pileup showing efficient deletion of exon 2 of *Sox9* in P14 retinas. **(d)** UMAP plot of combined P14 and P120 control and *Sox9*-deficient retinas. **(e)** UMAP plot of subsetted MG showing the clustering of P14 and P120 control and *Sox9*-deficient MG. **(f)** Violin plots of MG markers (*Glul*, *Abca8a*), Sox transcription factors (*Sox8*, *Sox5*, *Sox6*, *Sox2*), neurogenic transcription factor *Ascl1*, and anti-proliferative factor *Btg2*. **(g, g)** Heatmap of top 20 upregulated and downregulated genes in P14 **(g)** and P120 **(h)** *Sox9*-deficient retinas. **(i)** Gene set enrichment analysis of significant differentially expressed genes in *Sox9*-deficient MG compared to control MG at P14.

**Fig. S4.**
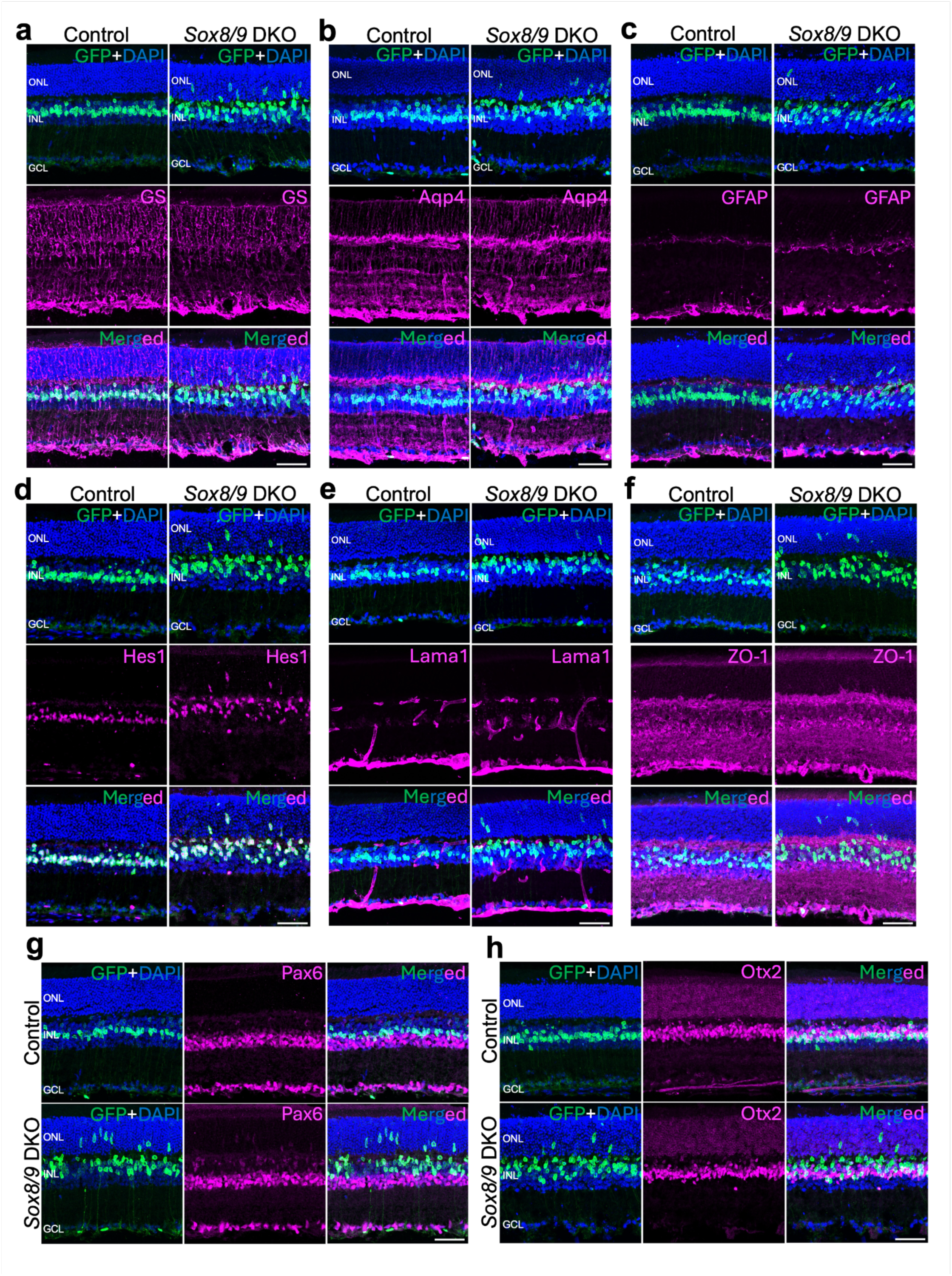
Immunohistochemical analysis of glial and neuronal markers in control and Sox8/9-deficient Müller glia. (a-d) Representative images of control and *Sox8/9* DKO retinas immunolabeled for GFP and glial markers, including **(a)** glutamine synthetase (GS), **(b)** Aqp4, **(c)** GFAP, and **(d)** Hes1. **(e-h)** Representative images of control and *Sox8/9* DKO retinas immunolabeled for GFP and **(e)** Lama1, **(f)** ZO-1, **(g)** Pax6, and **(h)** Otx2. ONL, outer nuclear layer; INL, inner nuclear layer; GCL, ganglion cell layer. DAPI, 4′,6-diamidino-2-phenylindole. Scale bar = 50µm.

**Fig. S5.**
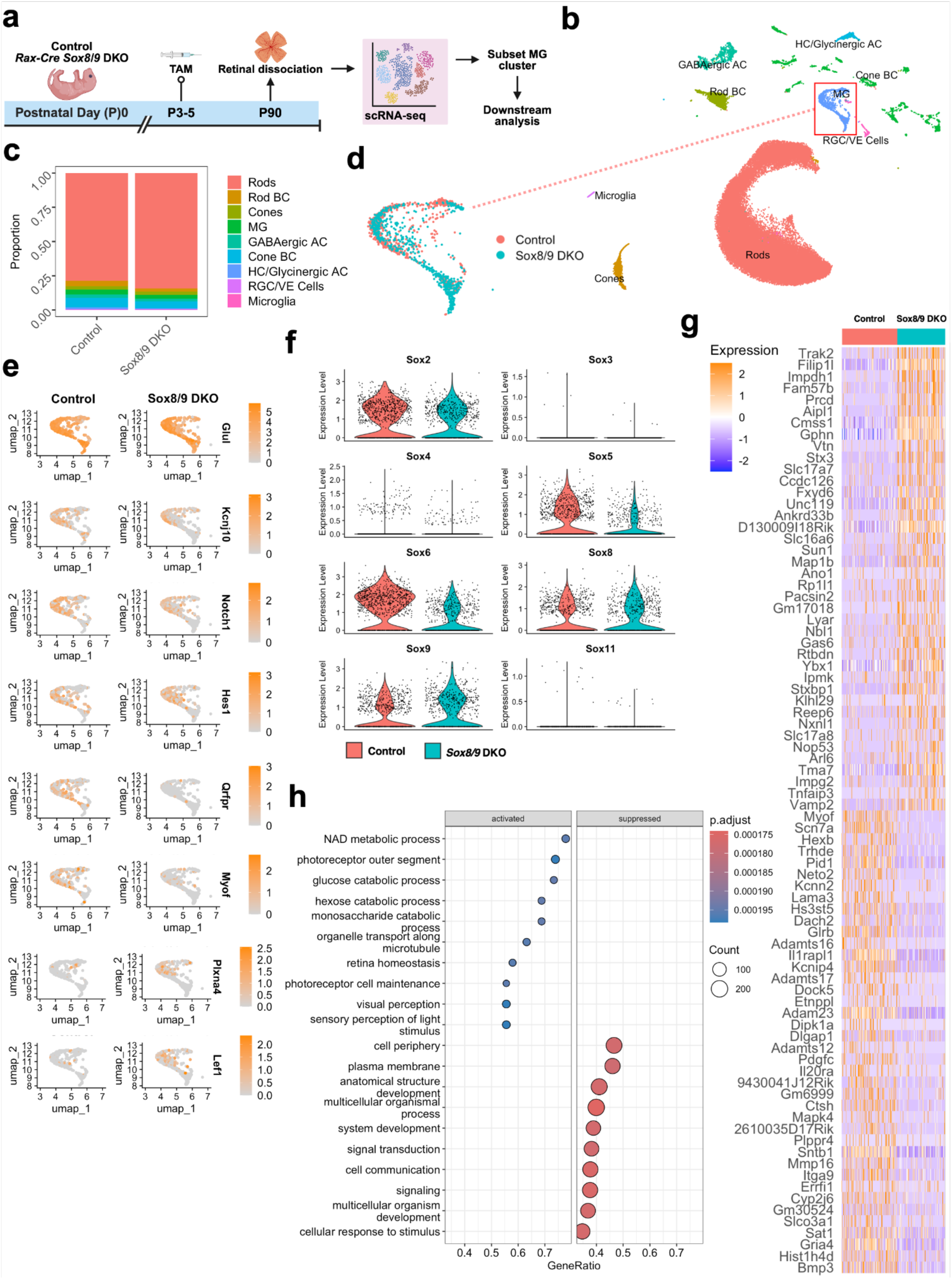
scRNA-seq analysis of control and *Sox8/9* DKO mature Müller glia. **(a)** Schematic of the scRNA-seq experimental pipeline. **(b)** UMAP plot of scRNA-seq datasets showing the clustering of all major retinal cell types in control and *Sox8/9*-deficient retinas. **(c)** Stacked bar plots representing the proportion of cells in each cluster between control and *Sox8/9*-deficient retinas. **(d)** UMAP plot of scRNA-seq datasets showing the clustering of all major retinal cell types in control and *Sox8/9*-deficient retinas. **(e)** Heatmap showing expression of the top 40 upregulated and downregulated genes in *Sox8/9*-deficient MG. **(f)** Feature plots highlighting differentially expressed genes associated with glial cell identity (*Glul*, *Kcnj10*) Notch signaling (*Notch1*, *Hes1*), cell migration (*Qrfpr*, *Myof*), cytoskeletal remodeling (Plxna4) and Wnt signaling (*Lef1*). **(g)** Violin plots showing decreased expression of Sox factors in *Sox8/9*-deficient MG. **(h)** Gene set enrichment analysis of significant differentially expressed genes in *Sox8/9*-deficient MG compared to control MG at P90.

**Fig. S6:**
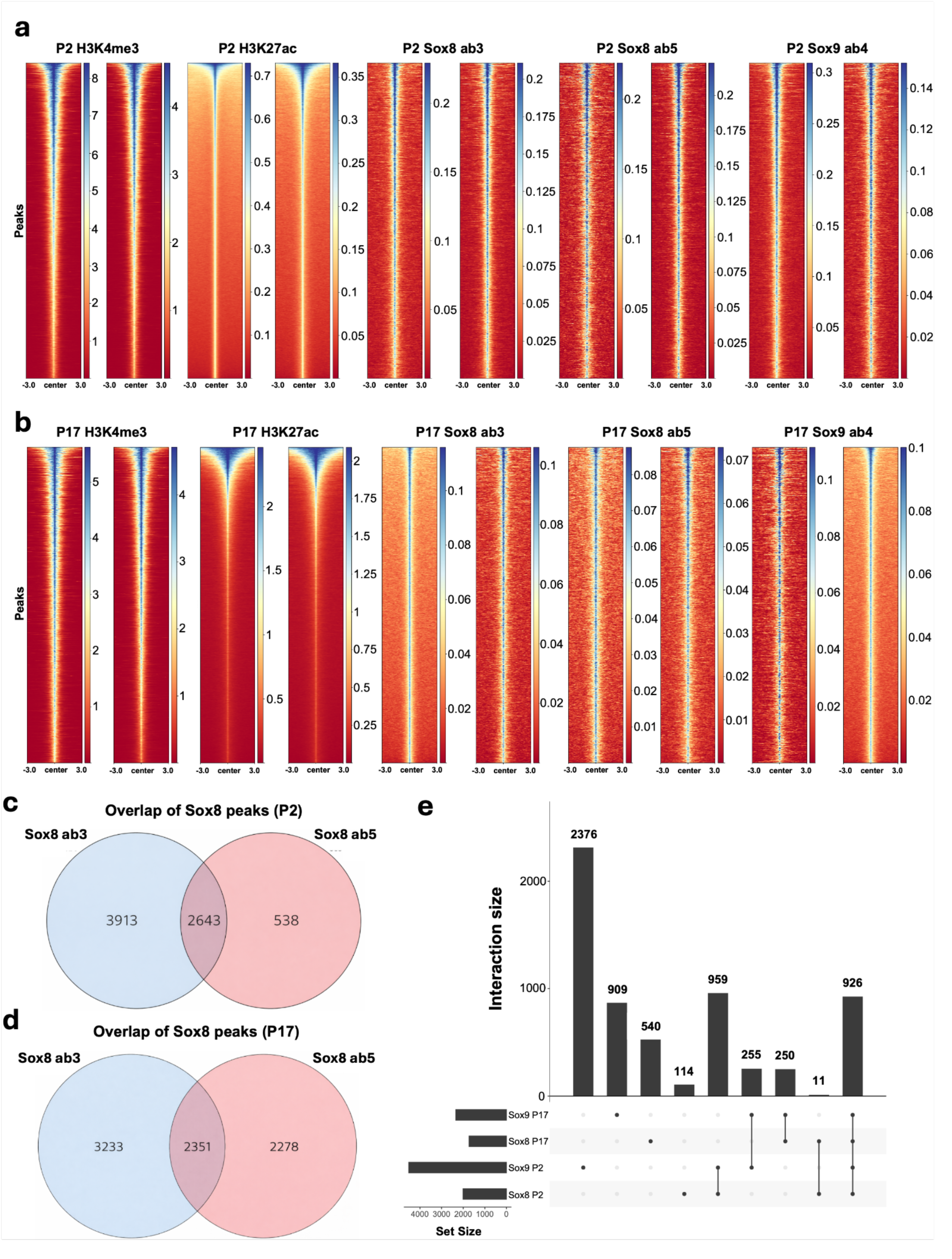
CUT&RUN analysis of Sox8 and Sox9: (a-b) Peak intensity plots for H3K4me3, H3K27ac, anti-Sox8 ab3, anti-Sox8 ab5 and anti-Sox9 ab4 at **(a)** P2 and **(b)** P17. Two replicate experiments were performed at both ages using each antibody. **(c-d)** Overlap of Sox8 peaks detected with ab3 and ab5 at **(c)** P2 and **(d)** P17. **(e)** Bar plot showing the number of overlapping peaks detected with Sox8 and Sox9 antibodies at P2 and P17.

**Fig. S7.**
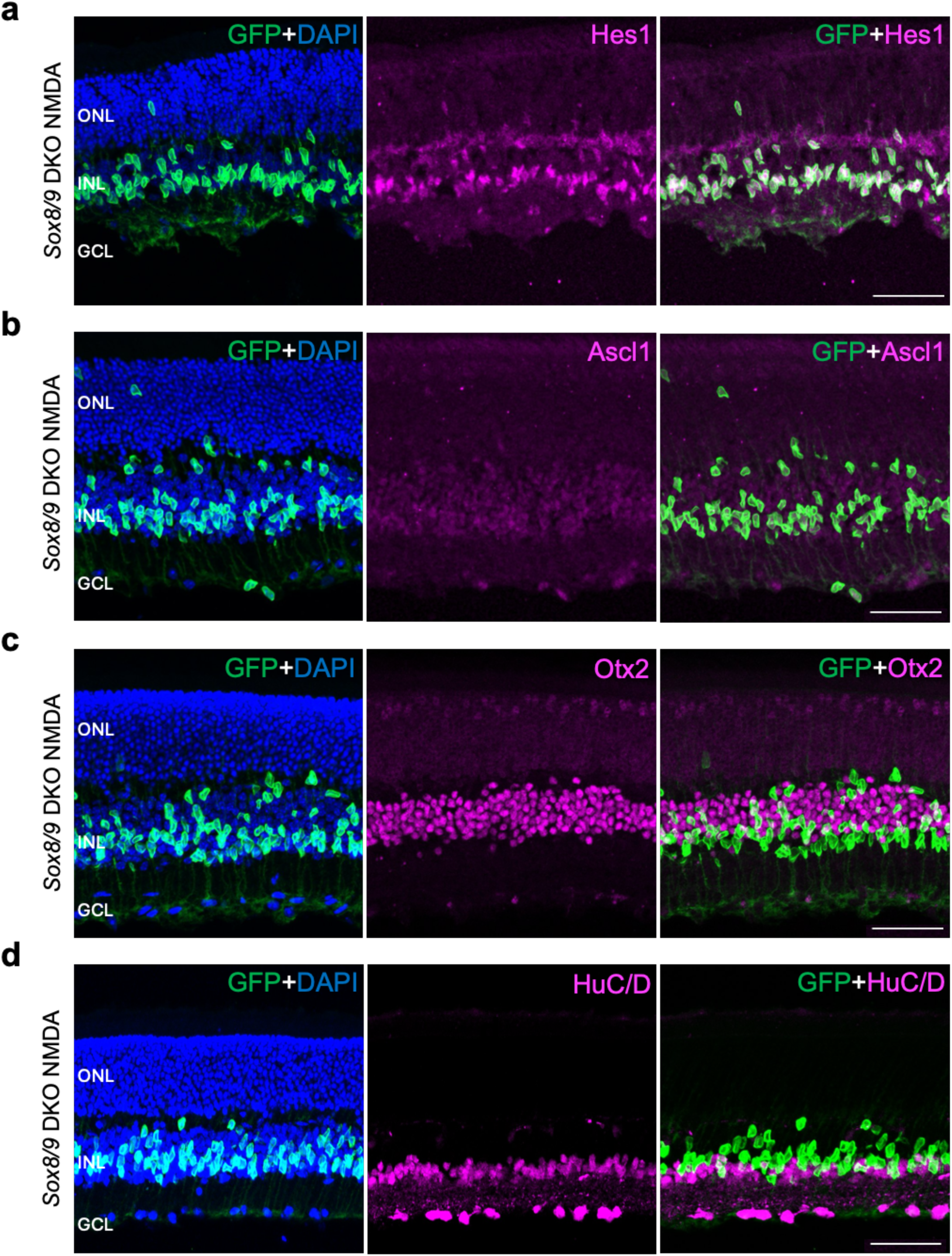
Selective loss of function of *Sox8/9* in mature MG does not induce neurogenesis with retinal injury. (a-d) Representative images of *Sox8/9* DKO retinas immunolabeled for glial **(a)** and neurogenic **(b-d)** markers, including **(a)** Hes1, **(b)** Ascl1, **(c)** Otx2, and **(d)** HuC/D. Scale bar = 50µm.

**Table ST1**: ScRNA-Seq analysis of P14 and P120 *Chx10-Cre;Sox9^lox/lox^*retina.

**Table ST2**: ScRNA/ATAC-Seq multiome data from *Glast-CreER;Sox8/9^lox/lox^* retina. Tamoxifen was administered via daily intraperitoneal injections at P3-5 and GFP-positive cells isolated via FACS at P17.

**Table ST3**: ScRNA-Seq data from *Rax-CreER;Sox8/9^lox/lox^* retina. Tamoxifen was administered via daily intraperitoneal injections at P3-5 and retinas harvested at P90.

**Table ST4**: CUT&RUN analysis of Sox8/9 and histone modifications in P2 and P17 retina

**Table ST5**: List of antibodies and commercial assays used in the study. Columns identify the reagent, source, catalog number, and RRID identifier if applicable.

**Table ST6**: Cell counts for immunohistochemical data.

## Methods

### Mice

Maintenance and experimental procedures performed on mice were in accordance with the protocol approved by the Institutional Animal Care and Use Committee (IACUC) at the Johns Hopkins School of Medicine under protocol number MO22M22. Mice were raised and housed in a climate-controlled pathogen-free facility on a 14-hour/10-hour light/dark cycle. *RaxCreER;Sun1-GFP* were used in this study and were generated by crossing the *RaxCreER* line developed by our group and the *Sun1-GFP* line developed by and obtained from J. Nathans at Johns Hopkins. *RaxCreER;Sox8/9^lox/lox^;Sun1-GFP* mice were generated by crossing *RaxCreER;Sun1GFP* mice with conditional *Sox8^lox.lox^*(Cambridge-Soochow University Genome Resource Center, EPD0618_3_E09 )and *Sox9^lox/lox^*(The Jackson Laboratory, RRID:IMSR_JAX:013106) mice. *GlastCreER;Sun1-GFP* mice were also used in this study and were generated by crossing the *GlastCreER* and *Sun1-GFP* lines developed by J. Nathans at Johns Hopkins and were obtained from his group. *GlastCreER; Sox8 ^lox/lox^;Sun1-GFP*, *GlastCreER;Sox9 ^lox/lox^;Sun1-GFP*, and *GlastCreER;Sox8/9 ^lox/lox^;Sun1-GFP* mice were generated by crossing Cre-negative *Sox8/9 ^lox/lox^;Sun1-GFP* mice with *GlastCreER;Sun1-GFP* mice. *Chx10Cre* (Rowan and Cepko, 2004) mice were crossed with *Sox9^lox/lox^* mice to generate *Chx10Cre;Sox9 ^lox/lox^* mice. Timed pregnant CD1 mice were ordered from Charles River Laboratories for the explant plasmid overexpression analysis.

### TAM injection

To induce Cre recombination, animals at postnatal day (P)3 were intramuscularly injected with TAM (Sigma-Aldrich, no. H6278-50mg) in corn oil (Sigma-Aldrich, no. C8267-500ML) at 0.5 mg per dose for three consecutive days.

### TAM diet

To induce Cre recombination, animals at ∼3 weeks of age were fed a TAM diet (Envigo, TD.130856) for three consecutive weeks.

### NMDA treatment

One week after finishing the TAM diet, adult mice were anesthetized with isoflurane inhalation. 1.5 microliters of 100 mM NMDA+1 μg/mL EdU in phosphate-buffered saline (PBS) were intravitreally injected using a microsyringe with a 33-gauge blunt-ended needle.

### EdU treatment

At the same time as NMDA treatment, mice were administered EdU (Abcam, no. ab146186) via intraperitoneal injections [150 μl of PBS (5 mg/ml)]. Additional intraperitoneal injections were performed daily for the following two days, totaling three injections over three days.

### Retinal Explant Electroporation and Culture

Eyes were enucleated from E14.5 CD1 embryos, and retinas were dissected in fresh, cold 1× PBS, then transferred to a micro electroporation chamber. The retinas were electroporated with plasmid solution at 30 V of 50-ms duration with 950-ms intervals (Matsuda and Cepko 2008) using either a pCAGIG plasmid that expresses IRES-GFP (Matsuda and Cepko 2004; Onishi et al. 2009), a bicistronic transcript that expresses full-length human *SOX8* or *SOX9* with IRES-GFP, or a 50:50 mixture of the *SOX8* and *SOX9* constructs (1ug/μl). The electroporated retinas were then flattened by a series of radial cuts and mounted on 0.2 μm Nucleopore Track-Etch membranes on culture media (For 50ml final media, 12.5ml of HBSS, 12.5 ml of heat-inactivated horse serum, 1.45ul of glucose solution, 25ml of MEM, 500ul of penicillin/streptomycin, and 50ul of L-glutamine was added) in 12-well plates and cultured until E19 (P0). At E16, retinal explants were pulsed with EdU (5mg/ml in PBS).

### Immunohistochemistry and imaging

Collection and immunohistochemical analysis of retinas were performed as described previously (de Melo, et al. 2016). Briefly, mouse eye globes and E19 (P0) retinal explants were fixed in 4% paraformaldehyde (Electron Microscopy Sciences, no. 15710). Eye globes were fixed for 4 hours at 4°C, while retinal explants were fixed for two hours at 4°C. Retinas were dissected and explants were carefully detached from the membrane in 1X PBS, followed by incubation in 30% sucrose overnight at 4°C. Retinas were then embedded in frozen section medium (VWR, no. 95057-838), cryosectioned at 16-μm thickness, and stored at −20°C. Sections were dried for 30 min in a 37°C incubator and washed three times for 5 min with 0.1% Triton X-100 in PBS (PBST). EdU labeling was performed by using a Click-iT EdU kit (Thermo Fisher Scientific, no. C10340) following the manufacturer’s instructions. Sections were then incubated in 10% horse serum (Thermo Fisher Scientific, no. 26050070), 0.2% Triton X-100 in 1X PBS (blocking buffer) for one hour at room temperature (RT) and then incubated with primary antibodies in the blocking buffer overnight at 4°C. Primary antibodies used are listed in Table ST5.

Sections were washed three times for 5 min with PBST to remove excess primary antibodies and were incubated in secondary antibodies in blocking buffer for one hour at RT. Sections were then washed three times for 5 min in PBST, once in PBSm and mounted with Fluoromount-G™, with DAPI (Thermo Fisher Scientific, no. 00-4959-52) under coverslips (VWR, no. 48404-453), air-dried, and stored at 4°C. Fluorescent images were captured using a Zeiss LSM 700 confocal microscope. Secondary antibodies used are listed in table ST5.

### Cell quantification and statistical analysis

Sox2/GFP-positive cells displaced to the ONL/OPL were counted and divided by the total number of Sox2/GFP-positive cells from a single random whole section per retina. EdU/GFP-positive cells were counted and divided by the total number of GFP-positive cells from a single random whole section per retina. Each data point in the bar graphs was calculated from an individual retina. All cell quantification data were graphed and analyzed using GraphPad Prism 10. Two-way analysis of variance (ANOVA) was used for analysis between three or more samples of multiple groups. All results are presented as means ± SD. Raw cell counts are listed in table ST6.

### Retinal cell dissociation

Retinas were dissected in fresh ice-cold PBS, and retinal cells were dissociated using an optimized protocol as previously described (Le et al. 2024). Each scRNA-seq sample contains two retinas from two animals of both sex. Each multiome sample contains a minimum of six retinas from six animals of mixed sex. Each CUT&RUN sample contains six retinas from three animals of mixed sex. Dissociated cells were resuspended in ice-cold Hibernate Buffer A GlutamaxBuffer containing Hibernate A (BrainBits, no. HALF500), B-27 supplement (Thermo Fisher Scientific, no. 17504044), and GlutaMAX (Thermo Fisher Scientific, no. 35050061).

### Single-cell RNA-seq library preparation

scRNA-seq was prepared on dissociated retinal cells using the 10x Genomics Chromium Single Cell 3′ Reagents Kit v3 or v4 (10x Genomics, Pleasanton, CA). Libraries were constructed following the manufacturer’s instructions and were sequenced using Illumina NovaSeq X. Sequencing data was processed through the Cell Ranger 8.0.1 pipeline (10x Genomics) using default parameters.

### Single-cell Multiome ATAC and GEX sequencing library preparation

scATAC-seq and scRNA-seq were prepared on FACS-isolated GFP-positive cells using the 10X Genomic Chromium Next GEM Single Cell Multiome ATAC and Gene Expression kit following the manufacturer’s instructions. Nuclei were isolated using an optimized protocol previously described(Santiago et al. 2023). Briefly, cells were spun down at 500*g* for 5 min and resuspended in 100 μl of ice-cold lysis Buffer. The cells were then lysed by pipette-mixing four times and incubated on ice for 4 min total. Nuclei were washed with 0.5 mL of ice-cold Wash Buffer and spun down at 500*g* for 5 min at 4°C. Nuclei pellets were resuspended in 15 μl of Nuclei Buffer and counted using DAPI. Resuspended nuclei (10,000 to 15,000) were used for transposition and loaded into the 10x Genomics Chromium Single Cell system. ATAC libraries were amplified with 7 PCR cycles and were sequenced on Illumina NovaSeq X with ∼400 million reads per library. RNA libraries were amplified from cDNA with 14 PCR cycles and were sequenced on the Illumina NovaSeq X platform.

### Single-cell RNA-seq data analysis

scRNA-seq data was analyzed using the Seurat R package(Stuart et al. 2021). Briefly, gene expression data were jointly normalized by function “NormalizeData”. The principal components analysis, dimensionality reduction, Louvain clustering, and UMAP visualization were performed on the top 15 principal components. Expression of known marker genes from previous studies were used to confirm the appropriate assignment of cell types (Lyu et al. 2021; Liu et al. 2023). Expression of significantly differentially expressed genes in MG was analyzed using the “FindMarkers” function. For multiomic analysis of control and *Sox8/9* double-knockout MG was performed using the Seurat and Signac workflow (Hao et al. 2024). Motif annotations were added using JASPAR2024 core vertebrate position frequency matrices, and transcription factor (TF) activity was quantified with chromVAR (Schep et al. 2017; Rauluseviciute et al. 2024). Differential gene expression (DEGs) and differential chromatin accessibility (DARs) between *Sox8/9* double-knockout and control MG were identified using Seurat’s FindMarkers with a logistic regression test for ATAC peaks using ATAC depth as a latent variable and standard Wilcoxon tests for RNA, applying an adjusted p < 0.05 threshold. DARs were annotated to nearest genes using ClosestFeature. Gene ontology enrichment of DEGs was performed using clusterProfiler (Yu et al. 2012).

### CUT&RUN library preparation and sequencing

C57BL/6J mice were euthanized and retinae were isolated in cold PBS containing 1X cOmplete™ EDTA-free Protease Inhibitor Cocktail (Millipore Sigma, 11873580001). Two biological replicates were collected from P2 and P17 mice, each containing a pool of 6 retinae from 3 mice. Retinas were collected into microcentrifuge tubes and snap frozen by submerging in an isopentane bath on liquid nitrogen for 1 minute and held at -80°C until nuclei extraction was performed. For nuclei extraction, the described protocol was followed, with minor modification (Tangeman et al. 2025). In brief, 1X cOmplete™ EDTA-free Protease Inhibitor Cocktail was added to all solutions, and lysis was performed by incubating tissue for 5 minutes in a lysis buffer containing 10 mM Tris-hydrochloride (pH 7.4) (Millipore Sigma, 93313-1L), 10 mM NaCl (Millipore Sigma, S5150-1L), 3 mM MgCl_2_ (Millipore Sigma, M1028), 0.01% Tween-20 (Bio-Rad, 1610781), 0.01% NP-40 Surfact-Amps™ Detergent Solution (ThermoFisher Scientific, 85124), 1% bovine serum albumin (Miltenyi Biotec, 130-091-376), and 1X cOmplete™ EDTA-free Protease Inhibitor Cocktail (Millipore Sigma, 11873580001), with gentle trituration throughout the incubation. Cold CUT&RUN wash buffer was prepared according to manufacturer’s guidance (EpiCypher, 14-1048) and was used as substitute for the final nuclei wash and resuspension. 600,000 nuclei were loaded per CUT&RUN reaction for use with Epicypher CUT&RUN protocol v5.1 and the CUTANA™ ChIC/CUT&RUN Kit (EpiCypher, 14-1048). A 1.8X SPRI bead ratio was used for the final recovery of CUT&RUN DNA.

The following antibody concentrations were used in the study: 0.5 µg/reaction IgG (EpiCypher, 13-0042K), 0.5 µg/reaction H3K4me3 (EpiCypher, 13-0060K), 0.5 µg/reaction H3K27ac (EpiCypher, 13-0059), 0.5 µg/reaction SOX8 (DSHB, DA11B7), 0.5 µg/reaction SOX8 (DSHB, PCRP-SOX8-2B9), 0.755 µg/reaction SOX8 (GeneTex, GTX129949), 0.5 µg/reaction SOX8 (DSHB, PCRP-SOX8-2D11), 0.6 µg/reaction SOX8 (Proteintech, 20627-1-AP), 0.5 µg/reaction SOX9 (DSHB, PCRP-SOX9-1A2), 0.5 µg/reaction SOX9 (DSHB, PCRP-SOX9-1E5), 0.618 µg/reaction SOX9 (Abcam, ab185966), and 0.5 µg/reaction SOX9 (Millipore Sigma, ab5535).

The SNAP-CUTANA™ K-MetStat Panel (EpiCypher, 19-1002k) was added to IgG, H3K4me3, and H3K27ac reactions. Final DNA was quantified using the Qubit 3 Fluorometer and the Qubit dsDNA HS Assay Kit (ThermoFisher Scientific, Q32854). Up to 5 ng of DNA per reaction was used for library preparation. The EpiCypher CUTANA™ CUT&RUN Library Prep protocol v1.5 was followed using the NEBNext® Ultra™ II DNA Library Prep Kit for Illumina® (NEB, E7645S). Provided adapters were diluted 1:25, per manufacturer’s recommendation. Indexing was performed usingNEBNext® Multiplex Oligos for Illumina® (Dual Index Primers Set 1) (NEB, E7600S) with 15 cycles of index amplification. Libraries were sequenced at the Novogene Sequencing Core (Sacramento, CA) on a 10B lane of the NovaSeq X Plus system (Illumina).

### CUT&Run data analysis

Raw FASTQ files were trimmed to remove adaptor sequences using the Trim Galore wrapper for cutadapt (Martin 2011). Trimmed reads were aligned to the *Mus musculus* mm10 and *E. coli* K-12 MG1655 spike-in reference genome using Bowtie2 (Langmead and Salzberg 2012). Duplicate reads and low-quality alignments were removed with Picard and Samtools prior to downstream analysis (Li et al. 2009; “Picard,” n.d.). Spike-in normalization was performed by scaling mouse-aligned read counts to the proportion of reads mapped to the *E. coli* genome. Peaks were called using SEACR in stringent mode with IgG controls used as background to identify high-confidence enriched regions (Meers et al. 2019). Motif enrichment analysis was performed using HOMER, which identified strong SOX motif enrichment specifically in CUT&RUN datasets generated with antibodies against SOX8 (ab3: GeneTex GTX129949; ab5: Proteintech 20627-1-AP) and SOX9 (ab4: Millipore Sigma ab5535)(Heinz et al. 2010). To generate a unified set of Sox8/9-bound regions, replicate peak files from SEACR were intersected for each antibody and age, and overlapping intervals were merged to obtain high-confidence Sox8 peak sets for P2 and P17. Sox8 and Sox9 peaks were then combined at each age by reducing overlapping intervals to create merged Sox8/9 peak sets. Combined Sox8/9 peak sets were intersected with developmental ATAC-seq DARs and Sox8/9 conditional knockout DARs in MGs identified previously to determine age-specific or chromatin accessibility changes associated with Sox binding. Peaks were annotated to genes and genomic features using ChIPseeker, with promoter, enhancer, and dual-marked status determined based on overlap with H3K4me3 and H3K27ac CUT&RUN peaks (Yu et al. 2015).

